# Rapid optogenetic blockade of autophagy reveals that nuclear pore complex proteins are robust autophagy substrates

**DOI:** 10.64898/2026.02.03.703609

**Authors:** Payel Mondal, Amanda Cyril, Dana Mamriev, Louis Parham, Igor Wierzbicki, Celina Shen, Ginevra Doglioni, Suzanne Dufrense, Anvita Komarla, David J. Gonzalez, Maximiliano A. D’Angelo, Christina G. Towers

## Abstract

Autophagy, a conserved recycling process, manages intracellular quality control to mitigate stress. To determine the rapid effects of autophagy perturbation, we developed the first optogenetic tool to rapidly inhibit autophagy, termed ASAP. Our approach leverages light-induced proteolytic cleavage to selectively inhibit autophagy within 5 minutes of light exposure, providing a precise and dynamic approach to study autophagy regulation. Proteomic profiling with ASAP revealed the most tightly regulated autophagy substrates along with novel, previously unidentified substrates, including nuclear pore complex (NPC) proteins. Interestingly, autophagy regulates quality control of cytoplasmic complexes of nucleoporins via specific LC3-interacting regions (LIRs), sparing nuclear pore complex proteins embedded in the nuclear envelope. Upon rapid autophagy inhibition, incomplete nucleoporin complexes accumulate and instead of undergoing autophagic degradation, cytoplasmic nucleoporin complexes aggregate in processing bodies (P bodies). Using ASAP, we demonstrate rapid and specific inhibition of autophagy, revealing that nuclear pore complex proteins are tightly regulated autophagy substrates.

## Main

Autophagy is a highly conserved cellular recycling process critical for maintaining cellular homeostasis^1, 2^. Accordingly, dysregulated autophagy is implicated across a wide spectrum of human diseases^3, 4^. Autophagy supports key developmental processes including the differentiation and maturation of various cell types^5^. Defective autophagy is associated with diverse diseases including neurodegeneration^6–8^, many types of cancer^9–12^, and broad pathologies associated with aging^13–15^. The dual nature of autophagy—as both a protective mechanism that maintains quality control in the basal state and during cellular stress, and as a contributor to disease when dysregulated—underscores its fundamental importance in human health and disease pathogenesis.

Autophagy regulates up to 70% of the proteome^16^. It degrades select proteins and organelles under both basal conditions and in response to stress. Selective mechanisms of autophagy that target organelles have been well characterized, including degradation of mitochondria (mitophagy), endoplasmic reticulum (ERphagy), and lipids (lipophagy), to name a few^17, 18^. While autophagy is presumed to regulate basal quality control of these organelles, most studies have been conducted in the context of specific organelle damage or nutrient stress. Beyond organelle turnover, autophagy also degrades proteins. While several studies have used unbiased techniques to identify autophagy substrates, including proteomics under conditions of stress and genetic deletion of core autophagy genes, various autophagosome isolation protocols, and in silico predictive studies based on sequence homology, there has been surprisingly little consensus among these assays^16, 19–21^. Autophagy is thought to maintain quality control under basal conditions; however, it has been difficult to assess these substrates in the absence of perturbations.

Under both basal conditions and during cell stress, autophagy is regulated by post-translational signaling within the cell to induce the formation of double membrane autophagosome structures^1^. This is mediated by upstream kinases that initiate membrane nucleation^22^. Subsequently, two ubiquitin-like conjugation systems facilitate the lipidation of ATG8 family proteins, including GABA type A Receptor-Associated Proteins (GABARAPs) and Microtubule-Associated Protein 1 Light Chain 3 (MAP1LC3)^23, 24^. For example, LC3-I is conjugated to phosphatidylethanolamine to form LC3-II, which is embedded in, and critical for, autophagosome membrane elongation and closure. Finally, soluble NSF (N-ethylmaleimide-sensitive fusion protein) attachment protein receptor (SNARE) proteins, such as syntaxin-17 (STX17), regulate the fusion of autophagosomes with lysosomes, ultimately leading to the degradation of the autophagosome contents and the membrane-embedded LC3-II^25–30^.

Autophagy responds to rapid post-translational signals, and in response, regulates cell fate rapidly. However, it has been difficult to study these immediate signaling events because the field lacked selective and rapid tools to inhibit autophagy. For example, lysosomotropic autophagy-targeting drugs lack specificity and accumulate within cells even after treatment is discontinued, preventing precise and reversible manipulation^31^. Genetic perturbations of key autophagy genes using either RNA silencing, CRISPR/Cas9, or doxycycline inducible systems are more specific but take days, if not weeks, to effectively block autophagy. Accordingly, the rapid effects of autophagy inhibition, including the most tightly regulated autophagy substrates have remained elusive.

Therefore, we created the first optogenetic tool to rapidly and reversibly inhibit autophagy. We demonstrate the utility of this tool to block autophagy within minutes with blue light and highlight its superiority to pharmacological autophagy inhibitors that target the lysosome. We leveraged this tool to identify the most tightly regulated autophagy substrates that accumulate immediately after autophagy inhibition. Under basal conditions, we found that outer mitochondrial membrane proteins are continuously turned over by autophagy, whereas mitochondrial matrix proteins are not. We also identified cytoplasmic nuclear pore complex (NPC) proteins, also known as nucleoporins, as bona fide autophagy substrates whose quality control is tightly regulated by select LC3-interacting regions (LIRs). Finally, we show that cytoplasmic nucleoporins that are not degraded by autophagy accumulate instead in processing bodies (P bodies), which form immediately after autophagy inhibition. Together, our studies highlight the utility of this optogenetic tool, termed ASAP, for rapidly inhibiting autophagy and identifying the most robust substrates, including NPC proteins. We predict this tool will have broad implications across the field of autophagy research.

## RESULTS

### Autophagic SNARE for Abrupt Photomanipulation (ASAP) inhibits autophagy within minutes

To create a tool to rapidly block autophagy, we leveraged a previously described truncation mutant of the key autophagy SNARE protein, STX17, which lacks the N-terminus, and acts as a dominant negative^30^. Previously, Uematsu et al. showed that 2 days of expression of myc-STX17ΔNTD blocks fusion of autophagosomes with lysosomes and inhibits autophagy, as evidenced by LC3-II accumulation. To leverage optogenetics to manipulate autophagy with light, we fused STX17ΔNTD to the photoactivatable protein hybrid LOV (hLOV1)^32^, which we engineered to cage a TEV protease recognition (tevS) site^33^ that is only accessible after a light-induced conformational change in hLOV1. To ensure equal expression, the TEV protease is transcribed from the same promoter and separated with a p2A consensus motif to enable ribosome skipping and allow for eukaryotic polycistronic RNA processing of both ASAP and TEV protease. The ASAP fusion construct is engineered such that it is targeted to the ER via an N-terminal IgK-leader sequence. Specifically, the construct utilizes this N terminal signal recognition sequence that is recognized through the signal recognition particle and localized to the signal recognition particle receptor on the ER. The construct includes a PDGFR single-pass transmembrane (TM)^34^ domain designed to anchor the protein to the endoplasmic reticulum (ER). In this configuration, the photoactivatable hLOV1 domain, the TEV protease cleavage site (tevS), and the effector DN-STX17 are presented on the cytoplasmic face of the membrane and will not be secreted. This design effectively tethers the entire construct to the ER in the dark, sequestering the dominant negative STX17 ΔNTD away from autophagosomes, allowing basal autophagy to proceed normally (Fig. 1a-b). Stimulation with pulsed 450nm light causes an immediate conformational change in hLOV1, uncaging the tevS, allowing cleavage of myc-STX17ΔNTD away from the ER such that it now binds to autophagosomes and prevents fusion with lysosomes. We coined this new optogenetic tool to inhibit autophagy <u>A</u>utophagic <u>S</u>NARE for <u>A</u>brupt <u>P</u>hotomanipulation (ASAP). Importantly, a control construct (CTRL) was also designed that includes a LOV domain fused to a tevS and Myc but does not include STX17ΔNTD or TEV and does not undergo cleavage to inhibit autophagy. Both ASAP and the CTRL are amenable to transient transfection in a variety of cultured cell lines.

**Figure 1.**
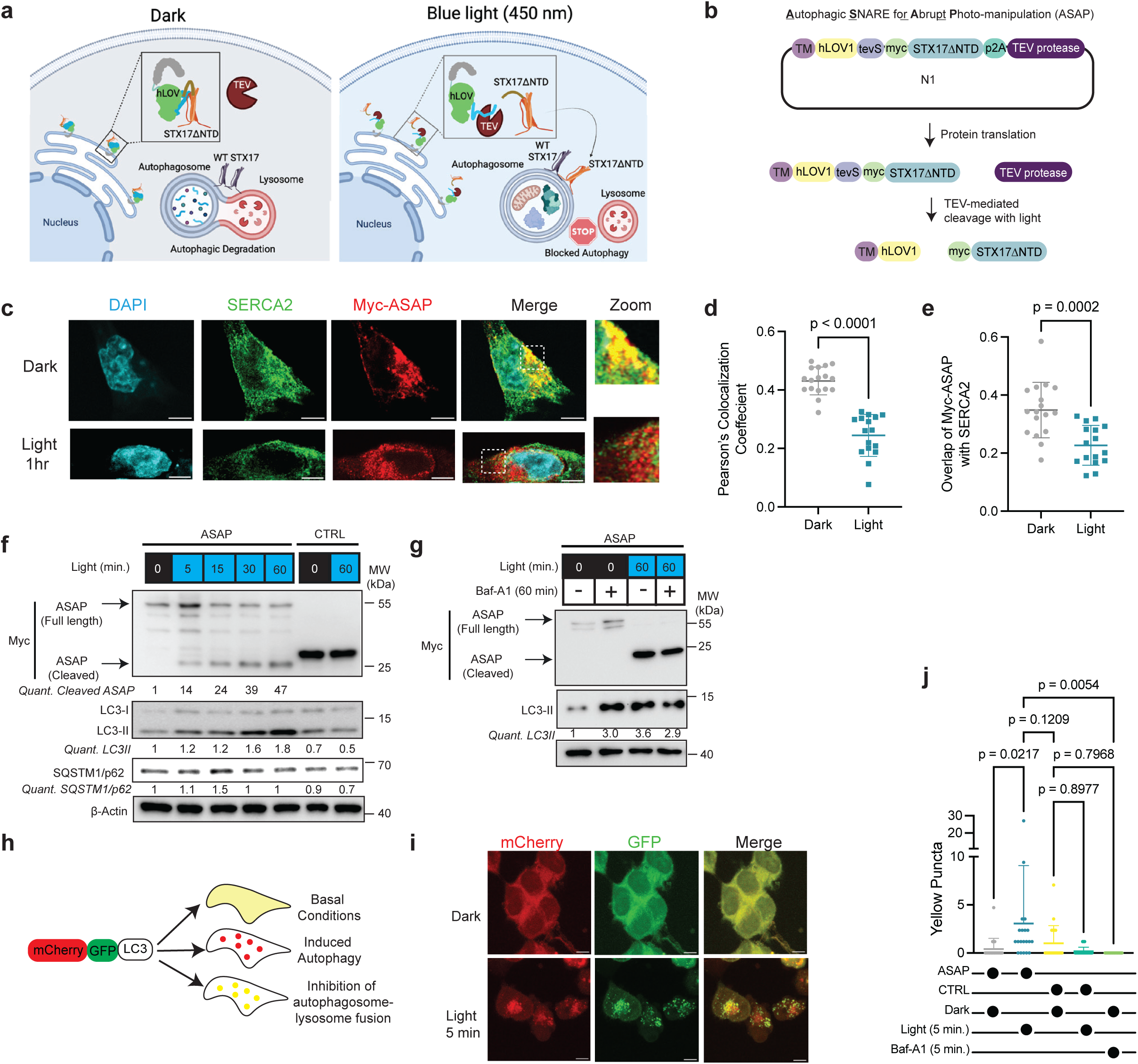
Autophagic SNARE for Abrupt Photomanipulation (ASAP) inhibits autophagy within minutes. **(a)** Model of the mechanism by which ASAP leads to autophagy inhibition in cells. **(b)** Schematic representation of the critical domains within the ASAP plasmid. **(c-e)** HEK293T cells transfected with ASAP and treated with 1hr of 450nm 10s on, 5 s off pulsed light. Immunofluorescence imaging (IF) of myc-ASAP (red) and the SERCA2 to mark the endoplasmic reticulum (ER; green). Quantification of **(d)** the Pearson’s colocalization coefficient between SERCA2 and Myc and **(e)** fraction of Myc overlapping with SERCA2. N=16-17 cells per condition. Data are graphed as mean ± S.D. and are representative of 3 independent experiments. Statistical analyses were performed using a two-tailed T-test. **(f)** HEK293T cells transfected with ASAP or CTRL and treated with indicated times of 450nm pulsed light. Western blot for Myc indicating full length ASAP, cleaved ASAP, and CTRL products, as well as LC3, SQSTM1/p62, and β-Actin. Quantification of the fold change in cleaved ASAP and LC3-II bands normalized to β-Actin are indicated. Blot is representative of N ≥ 3 experiments. **(g)** HEK293T cells transfected with ASAP and treated with 450nm pulsed light with or without bafilomycin (Baf-A1; 10nM) for 1hr to measure autophagy flux. Western blot for full length Myc, cleaved Myc, LC3, and β-Actin. Quantification of the fold change in LC3-II bands normalized to β-Actin are indicated. Blot is representative of N = 3 experiments. **(h)** Schematic representation of mCherry-GFP-LC3 autophagy reporter. **(i-j)** HCT116 cells with stable expression of mCherry-GFP-LC3 transfected with ASAP and treated with 450nm pulsed light for 5 minutes followed by IF for mCherry and GFP puncta. **(j)** Quantification of yellow LC3 puncta. N = 40-80 cells per condition. Data are represented as mean ± S.D for 2-4 independent experiments. Statistical analyses were performed using an Ordinary one-way ANOVA. Uncropped western blots are provided in Extended Data Fig. 9

Using different cell lines including HEK293T cells, HCT116 colon cancer, and NCI-H292 lung cancer cells, we first confirmed that ASAP inhibits autophagy. Immunofluorescence imaging and cell fractionation assays confirmed that Myc-tagged ASAP is colocalized with ER markers in the dark and treatment with pulsed blue light causes ASAP cleavage and dissociation from the ER (Fig. 1c-e, Extended Data Fig. 1a-c). Given the use of the TM domain, full length ASAP should be locked in the ER or potentially trafficked to the plasma membrane, however little ASAP was detected at the plasma membrane based on co-staining with the plasma membrane marker Wheat Germ Agglutinin (WGA) (Extended Data Fig. 1d). Immunoblotting for Myc after a rapid time course of blue light revealed an increase in cleaved ASAP starting as early as 5 minutes after blue light and progressively increasing with longer light exposures (Fig. 1f, Extended Data Fig. 1b-c). We observed a corresponding accumulation in LC3-II, indicative of STX17 inhibition of autophagosome-lysosome fusion. There is also a subtle and brief increase in the autophagy receptor protein p62/SQSTM1 within the first 15 minutes of light treatment. LC3 accumulation with light was confirmed by IF in Myc-ASAP expressing cells (Extended Data Fig. 1e-f). Importantly, even 60 minutes of light does not cause an increase in LC3-II or p62/SQSTM1 in cells expressing the CTRL construct (Fig. 1f, Extended Data Fig. 1c). To ensure that ASAP blocks autophagic flux, we included the lysosome inhibitor Bafilomycin-A1 (Baf-A1) in combination with light. Both 1hr of Baf-A1 alone or 1hr of light treatment alone in ASAP expressing cells causes an accumulation in LC3-II, however combining the two does not cause any further increase in LC3-II indicating that ASAP inhibits autophagy and does not increase autophagic flux (Fig. 1g, Extended Data Fig. 1b). Additionally, autophagic flux was measured using the pH sensitive mCherry-GFP-LC3 reporter, leveraging the pH stable fluorophore, mCherry, and pH sensitive fluorophore, GFP, which results in a mostly diffuse yellow signal under basal conditions^35^. Autophagosome-lysosome fusion events, or autolysosomes, present as red puncta, and accumulated autophagosomes that cannot fuse with lysosomes present as yellow puncta (Fig. 1h). A 4hr treatment of Baf-A1 is used as a positive control confirming an accumulation of yellow puncta with this assay (Extended Data Fig. 1g-h). Just 5 minutes of pulsed blue light in ASAP, but not CTRL, expressing cells causes a similar accumulation in yellow puncta (Fig. 1i-j), indicating robust and rapid inhibition of autophagosome-lysosome fusion. Live cell imaging revealed that CTRL cells stimulated with 450nm light via confocal microscopy exhibit transient yellow puncta that appear and then lose the GFP signal within 90 seconds on average. The stable mCherry signal and lack of GFP signal are prominent in these images indicating the presence of constant basal autophagy (Extended Data Fig. 2a-b). However, in ASAP-expressing cells stimulated with light, yellow puncta appear, indicating that the upstream autophagy machinery is still intact. These puncta do not resolve and persist for several minutes indicating the accumulation of autophagosomes that cannot fuse with lysosomes (Extended Data Fig. 2a-b).

Another version of ASAP was generated that is tagged with mRuby2 to allow for live cell imaging where ASAP transfected cells are identifiable with live cell fluorescent microscopy. Due to the bulky fluorescent protein, the mRuby2 is engineered on the N terminal side of the tevS and as a result even after light mediated cleavage mRuby2 remains on the ER membrane so that the bulky fluorescent protein does not hinder cleaved myc-STX17ΔNTD from translocating. Live cell imaging confirmed robust colocalization of mRuby2-ASAP with live cell ER tracker (Extended Data Fig. 2c). Myc cleavage and LC3 accumulation was observed with the mRuby2-ASAP construct as well (Extended Data Fig. 2d).

To further confirm minimal off target effects of the TEV protease, additional ASAP constructs were developed that do not contain either the tev site (ASAPΔtevs) or TEV protease (ASAPΔTEV). As expected, light does not induce any changes in LC3-II in cells expressing either ASAPΔtevs or ASAPΔTEV (Extended Data Fig. 2e). Interestingly, multiple high molecular weight bands near the predicted size for full length ASAP as well as ASAPΔtevs and ASAPΔTEV were observed, indicating these bands are not from off-target TEV cleavage but instead are likely light-independent degradation products.

### ASAP-mediated autophagy inhibition is reversible

To specifically benchmark ASAP against pharmacological autophagy inhibition, we compared 5 minutes of Baf-A1 treatment with 5 minutes of light in ASAP expressing cells. Baf-A1 fails to cause LC3 accumulation in CTRL cells treated with light for 5 minutes; however, 5 minutes of light alone in ASAP-expressing cells results in increased LC3-II (Fig. 2a).

**Figure 2:**
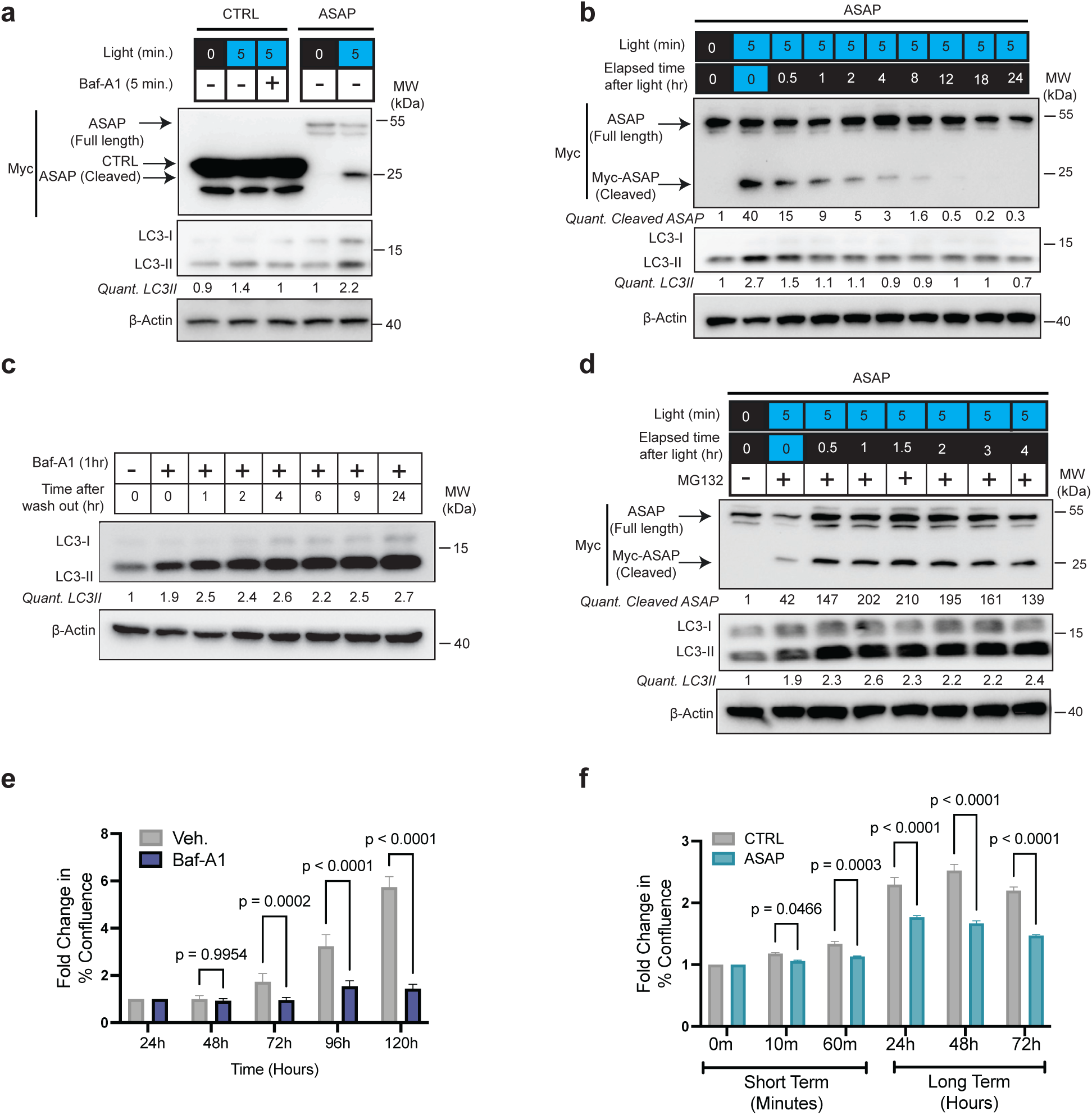
ASAP is reversible and blocks autophagy faster than pharmacological autophagy inhibition. **(a)** HEK293T cells transfected with ASAP or CTRL and treated with 450nm pulsed light for 5 minutes with or without Baf-A1 (10nM) for 5 minutes. Western blot for Myc, LC3, and β-Actin. Quantification in the fold change of LC3-II bands normalized to β-Actin are indicated. Blot is representative of N = 3 experiments. **(b)** HCT-116 cells transfected with ASAP and treated with 450nm pulsed light for 5 minutes after which the cells were returned to the dark and harvested after indicated periods of time. Western blot for Myc, LC3, and β-Actin. Quantification in the fold change of cleaved ASAP and LC3-II bands normalized to β-Actin are indicated. Blot is representative of N = 3 experiments. **(c)** HCT-116 cells treated with Baf-A1 (10 nM) for 1hr, then switched to media without Baf-A1 and harvested after indicated periods of time. Western blot for LC3 and β-Actin. Quantification in the fold change of LC3-II bands normalized to β-Actin are indicated. Blot is representative of N = 3 experiments. **(d)** HCT-116 cells transfected with ASAP and treated with 450nm pulsed light for 5 minutes after which the cells were returned to the dark and treated with MG132 (25 μM) then harvested after indicated periods of time. Western blot for Myc, LC3, and β-Actin. Quantification in the fold change of cleaved ASAP and LC3-II bands normalized to β-Actin are indicated. Blot is representative of N = 3 experiments. **(e)** HEK293 cells treated with Baf-A1 (10 nM) for 5 days, confluence analyzed by Incucyte live cell imaging every 24hrs. Data graphed as the fold change in % confluence normalized to time point 0 for each condition. Data are represented as mean ± S.D and are representative of 2 independent experiments. Statistical analyses were performed using a Two-way ANOVA and a Šídák’s multiple comparisons test. **(f)** HEK293 cells transfected with ASAP or CTRL and treated with pulsed blue light for short term time points, followed by pulsed light for 5 minutes every 30 minutes for 3 days for longer time points. Confluence analyzed by Incucyte live cell imaging at indicated time points. Data graphed as the fold change in % confluence normalized to time point 0 for each condition. Data are represented as mean ± S.D and are representative of 3 independent experiments. Statistical analyses were performed using a Two-way ANOVA and a Šídák’s multiple comparisons test. Uncropped western blots are provided in Extended Data Fig. 9

Next, we asked if ASAP-induced autophagy inhibition is reversible. ASAP-expressing cells treated with light for 5 minutes were then returned to the dark and harvested over time. Immunoblotting revealed that ASAP is cleaved after 5 minutes of light, but the cleaved protein quickly dissipates if light is removed resulting in a half-life of approximately 30 minutes (Fig. 2b). Accordingly, LC3-II accumulation resolves back to baseline within an hour of light removal. Baf-A1, however, takes 1-2 hours to induce LC3-II accumulation and removal of the drug is unable to reverse autophagy inhibition, even up to 24 hours after the drug was washed out (Fig. 2c). The reversibility of ASAP is prevented in the presence of the proteosome inhibitor, MG132, indicating that the mechanism of reversibility is because cleaved ASAP is unstable and quickly degraded by the proteosome (Fig. 2d).

Importantly, we confirmed that the intensity and duration of 450nm light required to activate ASAP are not detrimental to cells. Live cell imaging confirmed no detectable increase in caspase3/7 mediated apoptosis or ROS, nor is there a decrease in cell count in untransfected cells (Extended Data Fig. 2f-h). Since autophagy has been shown to support cell growth by providing critical nutrients and preventing apoptosis^36^, we compared the effects of pharmacological autophagy inhibition with Baf-A1 and ASAP on cell viability. While a significant growth defect was detected after 3 days of Baf-A1 treatment, ASAP expressing cells showed a growth defect within as early as 24hrs of pulsed blue light (Fig. 2e-f). However, there are minimal growth defects observed prior to 24hrs of light, suggesting that in HEK293 cells the detrimental effects of autophagy inhibition take longer to manifest.

Many optogenetics systems can leak resulting in TEV cleavage in the dark^37–39^. To test light-independent cleavage of ASAP, transfected cells were kept in the dark for either 16hrs (per the normal protocol), 24hrs, 40hr, or 48hrs prior to treatment with light. In all cases, cleaved ASAP is not detected, even 48hrs after transfection (Extended Data Fig. 2i). Importantly, autophagy inhibition, defined by LC3-II accumulation, was also observed at all time points with light. A slight increase in basal LC3-II expression was noted in the dark conditions at the later time points, and this is likely a result of cell stress due to overly crowded cells. Nonetheless, even at these time points, light causes an increase in LC3-II. Together these data indicate minimal light-independent TEV cleavage with the original ASAP system. Interestingly, the mRuby2-ASAP undergoes more robust TEV cleavage with the same amount of light. But as a consequence, more light-independent cleavage was also observed with the mRuby2-ASAP although the exact reason for this difference is unclear (Extended Data Fig. 2d). Together, these studies emphasize the utility of ASAP for rapidly and reversibly inhibiting autophagy, with superior time resolution compared to pharmacological agents targeting a similar step in the autophagy pathway.

### Proteomics profiling with ASAP reveals tightly regulated autophagy substrates

Autophagy is thought to maintain quality control under basal conditions; however, it has been difficult to assess these substrates in the absence of perturbations. While several studies have attempted to use unbiased proteomics to identify autophagy substrates, most of these studies have been in the context of different types of cell stressors, with little consensus on the top-regulated proteins. Therefore, we leveraged ASAP to block autophagy under basal conditions in the absence of stress to identify proteins that rapidly accumulate within 30 minutes, with the rationale that the proteins that accumulate the fastest after autophagy inhibition are the most tightly regulated and constitutively degraded autophagy substrates.

High-fidelity tandem mass tag (TMT) proteomics using MS3 mass spectrometry (MS) for quantification was performed in cells expressing CTRL or ASAP and treated with light for 5 or 30 minutes (Fig. 3a, Supplementary Table 1). Using Proteome Discover, we identified 5,219 high-confidence proteins representing all cellular compartments across our data sets. When comparing CTRL and ASAP cells both treated with light, there are far more proteins significantly increased after 30 minutes compared to 5 minutes as expected (Fig. 3b). To assess potential substrates, we focused on protein groups that start to trend to increase at 5 minutes and become significantly increased by 30 minutes. STX17 is one of the most increased proteins in this comparison due to its expression in the ASAP but not CTRL construct. Importantly, our data also confirms several previously identified autophagy substrates and adaptor proteins (Fig. 3c, Extended Data Fig. 3a)^16, 19–21^. While GABRAP is significantly increased, we do not see significant changes in all ATG8-family proteins. This is likely due to a limitation in proteomics that prevents it from accurately capturing small lipidated proteins and other membrane-embedded proteins^40^, as we confirmed that LC3B-II accumulates after 5 minutes of light in ASAP-expressing cells via western blot. Therefore, we acknowledge that a subset of small proteins cannot be assessed in this assay.

**Figure 3:**
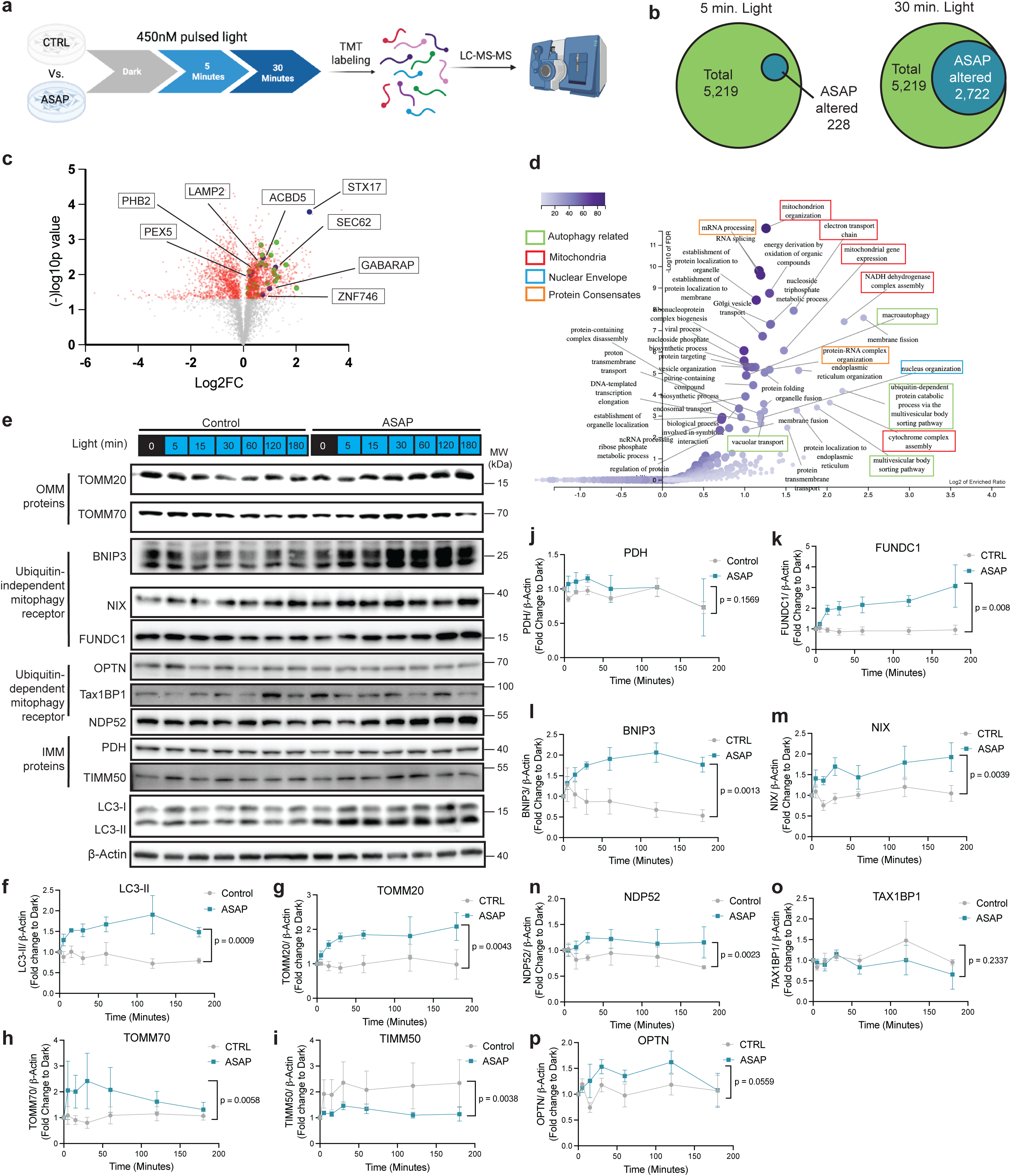
TMT Proteomics with ASAP reveals tightly regulated autophagy substrates. **(a)** Schematic overview of the tandem mass tag (TMT)-based quantitative proteomics workflow used to analyze protein abundance changes in HEK293T cells transfected with ASAP or CTRL and treated with 450nm pulsed light for 5 or 30 minutes. **(b)** Venn diagram showing the overlap between all proteins identified and those with statistically significant abundance changes in ASAP expressing cells relative to CTRL cells after 5 or 30 minutes of 450nm pulsed light. **(c)** Volcano plot depicting Log_2_ fold change and (-)Log_10_ P value for proteins altered in ASAP expressing cells versus CTRL cells after 30 minutes of 450nm pulsed light. Proteins with a p value < 0.05 are indicated in red. ATG8 family proteins, STX17 and known autophagy substrates are highlighted in blue and green. **(d)** Overrepresentation analysis (ORA) from proteins identified as significantly increased in ASAP compared to CTRL expressing cells after 30 minutes of light. Volcano plot using the enrichment category: Gene Ontology Biological Process non-redundant. **(e)** HEK293T cells transfected with ASAP or CTRL and treated with 450nm pulsed light for indicated periods of time. Western blot for mitochondrial proteins located in mitochondrial matrix, outer mitochondrial membrane, canonical mitophagy receptors, ubiquitin-independent mitophagy receptors, along with Myc, LC3, and β-Actin. Blot is representative of N ≥ 3 experiments. **(f-p)** Quantification of western blot bands normalized to β-Actin are indicated for indicated proteins graphed as the fold change to the respective dark conditions for ASAP and CTRL expressing cells. Data shown as the average ± SEM for N ≥ 3 experiments. Statistical analyses were performed using data from all timepoints and an unpaired two-tailed T-test with Welch’s Correction between ASAP and CTRL. Uncropped western blots are provided in Extended Data Fig. 9

We compared our data set of putative autophagy substrates, i.e. proteins that accumulate after 30 minutes of light, with previously published MS data sets that used various conditions to identify autophagy substrates. Prior studies used a combination of stable genetic perturbations of different autophagy genes across different model systems and stress contexts. Although there is some overlap with each data set, many proteins in our data remain unique and were not detected in these other studies (Extended Data Fig. 3b-e)^16, 19–21^.

Gene ontology analyses of accumulating proteins revealed numerous biological processes (Fig. 3d). Importantly, pathways related to autophagy and many selective autophagic degradation pathways were identified. Proteins related to mitochondria are also amongst the most significantly altered (Fig. 3d, Extended Data Fig. 3f). Mitochondria – via the selective autophagy process of mitophagy – are arguably the most well-characterized autophagy substrate^41^. Many studies have elucidated how a variety of mitochondrial insults induce mitophagy through different mechanisms including ubiquitin-dependent processes that require PINK1 and Parkin as well as ubiquitin-independent processes that rely on different mitophagy resident autophagy receptors^42–45^. Our proteomics results confirm that mitochondria are among the most robust autophagy substrates, even under basal conditions without mitochondrial perturbations.

To focus on specific mitochondrial proteins, we performed individual western blots for proteins representing different mitochondrial compartments using an acute time course of light treatment ranging from 5 minutes to 3 hours of autophagy inhibition (Fig. 3e-p). Like LC3-II, there is an immediate and sustained increase in proteins localized to the outer mitochondrial membrane (Fig. 3e-h). However, proteins localized to the mitochondrial matrix do not accumulate in this short time course (Fig. 3e, 3i-j). Interestingly, outer-mitochondrial membrane-resident, ubiquitin-independent mitophagy receptors like BNIP3, BNIP3L/NIX, and FUNDC1 accumulate rapidly with light-mediated autophagy inhibition while the respective mRNA expression remains stable (Fig. 3e, 3k-m, Extended Data Fig. 3g-j). However, mitophagy receptors involved in the PINK1/Parkin pathway, like OPTN, TAX1BP1, and NPD52, are less likely to accumulate at these early time points (Fig. 3e, 3n-p). Together, these results suggest that selective mitophagy mechanisms are engaged under basal conditions, and bulk mitochondria are not constantly degraded in line with the view that mitophagy regulates mitochondrial quality control without degrading mitochondria in bulk. To determine if there are rapid changes in mitochondrial dynamics immediately after autophagy inhibition, we performed immunofluorescence staining (IF) for mitochondrial proteins. The mitochondria appear more hyperfused with fewer individually counted mitochondria and longer branch lengths per mitochondria indicating an immediate mitochondrial stress response with the most apparent changes observed within 1-3hrs after autophagy inhibition (Extended Data Fig. 3k-m). Similar effects on mitochondria count and branch length were observed with sustained autophagy inhibition up to 48hrs with pulsed light in ASAP expressing cells or with Baf-A1 treatment (Extended Data Fig. 3n-q). Together these results emphasize that our proteomics strategy to utilize light-mediated rapid autophagy inhibition accurately identifies autophagy substrates.

### Cytoplasmic nucleoporins accumulate after rapid autophagy inhibition

Confident in our proteomics approach, we then aimed to discover previously unidentified autophagy substrates. One of the most significantly altered pathways from the gene ontology analyses is related to nuclear organization and the nuclear envelope (Fig. 3d). We identified several nuclear pore complex (NPC) proteins amongst the proteins that significantly accumulate after 30 minutes of light (Fig. 4a). The NPC is a highly organized protein complex that consists of dozens of different nucleoporins which together regulate the export and import of various proteins and mRNAs between the nucleus and cytoplasm^46^. Prior studies have shown that some NPC proteins, especially those localized to the inner ring complex, are among the most stable proteins in the proteome; however, other NPC proteins are less stable^47–49^. Studies in yeast indicate that some NPCs can be targeted to autophagosomes, although the exact mechanisms and their conservation in mammalian systems remain unknown^50–52^. Our studies represent the first indication in mammalian cells that NPC complex proteins could be targeted by autophagy.

**Figure 4:**
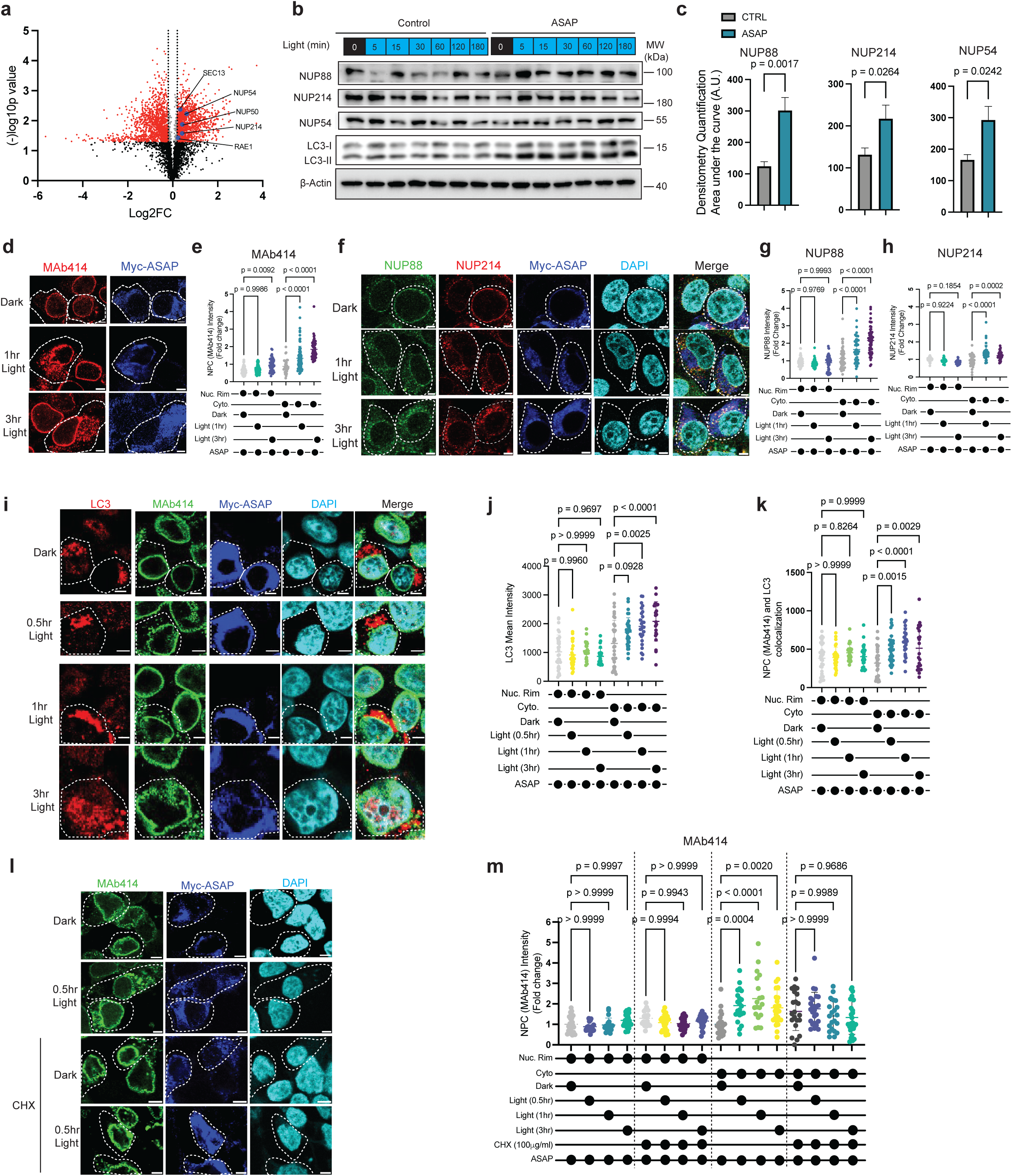
Cytoplasmic nuclear pore complex proteins accumulate after rapid autophagy inhibition. **(a)** Volcano plot from proteomics performed in HEK293T cells depicting Log_2_ fold change and Log_10_ P value for proteins altered in ASAP expressing cells versus CTRL cells after 30 minutes of 450nm pulsed light. Proteins with a p value < 0.05 and Log_2_ FC >0.2 are indicated in red and are the same as those depicted in Fig. 2C. Nuclear pore complex (NPC) proteins are highlighted in blue. **(b-c)** HEK293T cells transfected with ASAP or CTRL and treated with 450nm pulsed light for indicated periods of time. (**b**) Western blot for NUP88, NUP214, NUP54, LC3 and β-Actin. Blot is representative of N = 3 experiments. Note: These same lysates were ran for NUPs here and mitochondrial proteins so the same LC3 blot is shown in Fig. 3e and Fig. 4b. **(c)** Quantification of western blot bands for indicated proteins normalized to β-Actin are indicated and graphed as the fold change to the respective dark conditions for ASAP and CTRL expressing cells based on the area under the curve across the time course. Full curves for this data are shown in Extended Data Fig. 4a-c. Data shown as the average ± SEM for N ≥ 3 experiments. Statistical analysis was performed on the area under the curve (AUC) followed by two-tailed T-test between ASAP and CTRL. **(d-h)** HEK293T cells transfected with ASAP and treated with 450nm pulsed light for indicated periods of time followed by IF. **(d,f)** Representative IF images of the pan NPC antibody (MAb414), NUP88, NUP214 and Myc-tag for ASAP, as indicated. Dotted white lines indicate examples of ASAP positive cells. Scale bar = 5μM. **(e,g,h)** Quantification in the nucleoporin signal that is cytoplasmic or on the nuclear rim, data graphed as fold change to the respective dark condition for cytoplasmic and rim signal. N=40-80 cells per condition. Data are graphed as mean ± S.D. and are representative of 3 independent experiments. Statistical analyses were performed using a One Way ANOVA. **(i-k)** HEK293T cells transfected with ASAP and treated with 450nm pulsed light for indicated periods of time followed by IF. **(i)** Representative IF images of MAb414, Myc-ASAP, LC3, and Dapi. Dotted white lines indicate examples of ASAP positive cells. Scale bar = 5μM. (**j**) Quantification of LC3 intensity reported as raw pixel intensity. (**k**) Quantification of MAb414 colocalization with LC3 on the nuclear rim or in the cytoplasm, reported as the pixel intensity of the overlapping signal. N = 23-38 cells per condition. Data are graphed as mean ± S.D. and are representative of 2 independent experiments. Statistical analyses were performed using a One Way ANOVA. (**l-m**) HEK293T cells transfected with ASAP and treated with 450nm pulsed light for indicated periods of time with or without CHX (10μg/ml) for the same amount of time (Dark samples were treated with CHX for 30 minutes) followed by IF. **(l)** Representative IF images of MAb414, and Myc-ASAP. Dotted white lines indicate examples of ASAP positive cells. Scale bar = 5μM. **(m)** Quantification of MAb414 on the nuclear rim or in the cytoplasm. Data graphed as fold change to the respective dark condition for cytoplasmic and rim signal. N = 19-25 cells per condition. Data are graphed as mean ± S.D. and are representative of 2 independent experiments. Statistical analyses were performed using a One Way ANOVA. Uncropped western blots are provided in Extended Data Fig. 9

Immunoblotting for individual NPC proteins confirmed rapid and sustained protein accumulation of NUP54, NUP88, and NUP214 in whole-cell lysates as early as 5 minutes after light-induced autophagy inhibition (Fig. 4b-c, Extended Data Fig. 4a-c). There is no increase in the transcript levels during the same time course indicating the increased protein expression is likely due to altered turnover versus increased gene expression (Extended Data Fig. 4d-f). While we identified nucleoporins as autophagy substrates after rapid autophagy inhibition, pharmacological inhibition also leads to nucleoporin accumulation (Extended Data Fig. 4g). Active NPCs are localized to the nuclear envelope; however, cytoplasmic NPC proteins have also been observed and can form either cytoplasmic aggregates or annulate lamellae (AL), which consist of many NPC proteins – although they lack specific proteins, making them incomplete NPCs – and are instead localized to the endoplasmic reticulum^53–56^. These cytoplasmic nucleoporins act as reservoirs for fully functional NPCs on the nuclear envelope^55, 57^. IF using a pan NPC antibody that detects multiple nucleoporins, as well as antibodies specific to NUP88, NUP214, and NUP93 revealed an accumulation in cytoplasmic NPC proteins immediately after autophagy inhibition with light in ASAP expressing cells; however NPC proteins localized to the nuclear rim remain largely unchanged (Fig. 4d-h, Extended Data Fig. 4h-k). Importantly there are no light-induced effects on cytoplasmic nucleoporins in cells expressing CTRL or ASAPΔtevS constructs (Extended Data Fig. 4l-o). Time course studies showed cytoplasmic accumulation of nucleoporins as early as 10 minutes after light, with peak accumulation after 3hrs; which is sustained with long term pulsed light out to 48hrs (Extended Data Fig. 4p). Short-term and long-term pharmacological autophagy inhibition leads to a similar, but slower, result (Extended Data Fig. 4q). Moreover, the accumulating NPC proteins colocalize with the increased LC3 in the cytoplasm, but not at the nuclear rim, to a greater extent in the ASAP expressing cells compared to CTRLs (Fig. 4i-k, Extended Data Fig. 5a-c). An interesting exception is, NUP153, which shows minimal cytoplasmic accumulation after light-mediated autophagy inhibition, and instead accumulates on the nuclear rim (Extended Data Fig. 4r-s). To determine if the accumulating cytoplasmic nucleoporins are derived from newly synthesized proteins or are sourced from existing NPCs, ASAP-expressing cells were treated with cycloheximide (CHX) to block protein translation followed by a time course of light treatment and IF for nucleoporins. As expected, cytoplasmic nucleoporins accumulate with light, while nucleoporins located on the nuclear rim are not affected. Interestingly, the effect on the cytoplasmic nucleoporins is blocked in the presence of CHX. This result indicates that the accumulating nucleoporins arise from newly synthesized nucleoporins and are not derived from pre-existing NPCs or free nucleoporins (Fig. 4l-m).

To confirm that cytoplasmic nucleoporin accumulation is not causing increased function of nuclear import through NPCs, Fluorescence Recovery After Photobleaching (FRAP) was performed using a GFP containing nuclear export and import sequences^58^. Light treatment in ASAP-expressing cells resulted in equal recovery times, indicating no change in NPC function at the nuclear envelope with 3hrs or even up to 72 hours of light treatment (Extended Data Fig. 5d-f). This supports our model that autophagy regulates NPC quality control by degrading cytoplasmic NPC protein reservoirs, sparing functional NPCs on the nuclear envelope.

### NUP214 and NUP88 have bona fide LC3 interacting regions

Next, we determined the mechanism underlying autophagic degradation of nucleoporins. The core sequence motif, [W/F/Y]_0_-X_1_-X_2_-[L/I/V]_3_, defines a canonical LIR^59–63^. NUP214 contains two predicted LIRs and NUP88 contains one predicted LIR. The predicted LIR1 sequence in NUP214 has strong conservation across vertebrate orthologs including mammals, monotremes, and birds, with preservation of the canonical aromatic and hydrophobic residues required for LC3 interaction. In contrast LIR2 is only conserved in primates. Similar to LIR1 in NUP214, the single LIR in NUP88 is highly conserved across vertebrates, excluding birds, suggesting these predicted LIR sequences are functional (Fig. 5a). Prior studies have shown that N<u>UP1</u>59 in yeast has a LIR and is a homologue of human NUP214^50–52^. The NUP214 predicted LIR motif is not sequence-homologous to the yeast Nup159 Atg8-interacting region, although both proteins contain LIR/AIM-like motifs consistent with potential convergent acquisition of autophagy-related short linear motifs.

**Figure 5:**
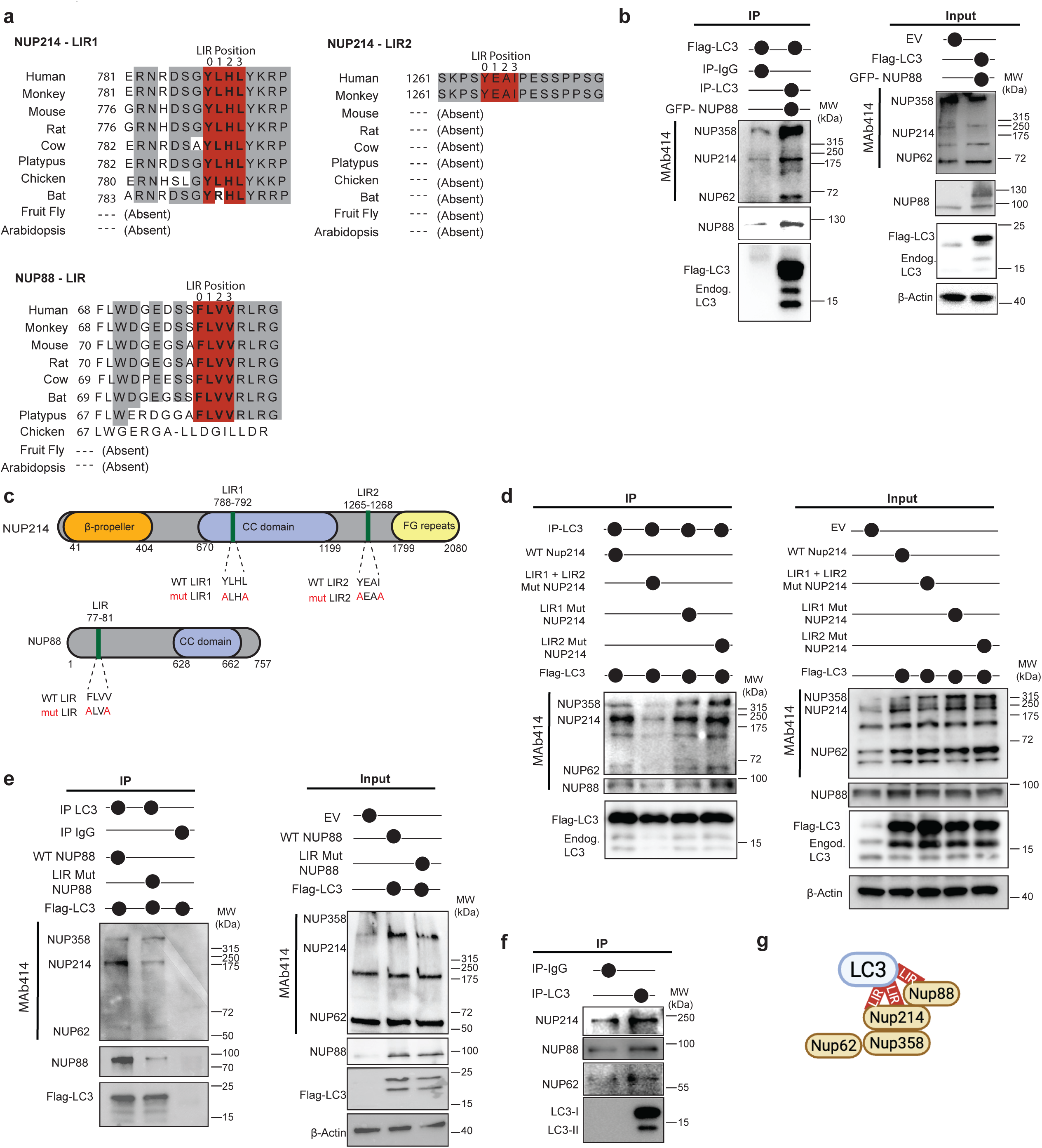
NUP214 and NUP88 have bona fide LC3 interacting regions. **(a)** Conserved sequence homology for the predicted NUP214 LC3 interacting region (LIR) 1, LIR2 and NUP88 LIR. **(b)** HEK293T cells expressing Flag-LC3 and GFP-NUP88 or PCDNA3.1 (EV). Immunoprecipitation pull down (IP) of LC3 followed by immunoblotting for MAb414 and LC3. (Left) Blots from IP elution and (right) blots from the input samples prior to IP. Blots are representative of 3 independent experiments. **(c)** Schematic of the (top) NUP214 and (bottom) NUP88 protein domains including the predicted wild type LIR sequences and the experimental mutations created within each LIR. **(d)** HEK293T cells expressing Flag-LC3 and either WT GFP-NUP214 or GFP-NUP214 with LIR mutants or PCDNA3.1 (EV). IP of LC3 followed by immunoblotting for MAb414 and LC3. (Left) Blots from IP elution and (right) blots from the input samples prior to IP. Blots are representative of 3 independent experiments. **(e)** HEK293T cells expressing Flag-LC3 and either WT or NUP88 mutants or PCDNA3.1 (EV). IP of LC3 followed by immunoblotting for MAb414 and LC3. (Left) Blots from IP elution and (right) blots from the input samples prior to IP. Blots are representative of 3 independent experiments **(f)** Endogenous IP of LC3 in HEK293T cells, followed by immunoblotting for MAb414 and NUP88. Blots are representative of 3 independent experiments. **(g)** Schematic representation of the suggested model by which LC3 interacts with the nuclear pore complex proteins. Uncropped western blots are provided in Extended Data Fig. 9

Immunoprecipitation (IP) of flag-LC3 followed by immunoblotting with the MAb414 antibody which detects several NPC proteins revealed that NUP214, NUP358, and NUP62 interact with LC3 (Fig. 5b). NUP88 was also detected in the immunoprecipitated fraction (Fig. 5b). To determine if these are bona fide LIRs that facilitate LC3 interaction and autophagic degradation of the nucleoporins, the key aromatic and aliphatic residues in the 0 and 3 positions were mutated to alanine (Fig. 5c). IPs of LC3 in the presence of WT or mutant GFP-NUP214 revealed that the interaction is dependent upon both LIRs in NUP214 as mutation of each alone does not disrupt the interaction but mutation of both LIRs together prevents NUP214 from immunoprecipitating with LC3 (Fig. 5d). Similarly, in the reverse IP, flag-LC3 is detected when WT NUP214 is immunoprecipitated, but the interaction is most disrupted when both NUP214 LIR sites are mutated (Extended Data Fig. 5g). These results also show that mutating NUP214 LIRs does not affect NUP214 binding to NUP358, NUP62, or NUP88. The residual LC3 that pulls down with NUP214 LIR 1+2 mutant could be because NUP88 has its own LIR and the NUP214 interaction with NUP88 is still intact. Therefore, the LIR in NUP88 was also assessed. IP of LC3 revealed that WT NUP88 immunoprecipitated with LC3, whereas NUP88 with LIR mutations disrupted this interaction, although some mutant NUP88 still interacts with LC3 (Fig. 5e). Likewise, the reverse IP shows that pull-down with mutant NUP88 results in less immunoprecipitated LC3 (Extended Data Fig. 5h). Importantly, IPs assessing endogenous LC3 also reveal an interaction with both endogenous NUP214 and NUP88 (Fig. 5f, Extended Data Fig. 5i).

To determine if the LIRs in NUP214 or NUP88 act as autophagy receptors to facilitate the interaction of LC3 with a larger complex of cytoplasmic nucleoporin proteins, we also assessed expression of additional NPC proteins after immunoprecipitation of LC3 in cells expressing NUP214 or NUP88 mutant LIR proteins. Interestingly, disruption of the LIRs in NUP214 prevents the interaction of the non-LIR containing nucleoporins including NUP358 and NUP62 with LC3 (Fig. 5d), while disruption of the NUP88 LIR still allows some NUP214, NUP358, and NUP62 to interact with LC3. Together these studies indicate that NUP214 acts as the more potent autophagy receptor, via both LIRs, to target cytoplasmic complexes of nucleoporins for autophagic degradation (Fig. 5g).

### Nucleoporins accumulate in P bodies after autophagy inhibition via NUP358

Proteomics also revealed an increase in proteins related to protein-RNA interactions, which often form various types of condensates in the cytoplasm, including processing bodies (P bodies) and stress granules (SGs) (Fig. 3d). Both P bodies and SGs are known to form rapidly in response to stress which we confirm using a short time course of puromycin and sodium arsenite followed by IF for DCP1A and G3BP, respectively (Extended Data Fig. 6a-f). To determine whether rapid autophagy inhibition affects these condensates, IF was performed in multiple cell lines expressing ASAP and CTRL to assess DCP1A and G3BP puncta formation. Interestingly, within 10 minutes of autophagy inhibition an immediate increase in the number of DCP1A puncta, indicative of P bodies, is observed in ASAP expressing cells, but not cells expressing CTRL or ASAPΔtevS constructs (Fig. 6a-b, Extended Data Fig. 6g-i). We found that light causes a robust increase in the number of DCP1A puncta in multiple cell lines but does not affect P body size (Extended Data Fig. 6j-q). Broad effects on all condensates were not observed, as there are no G3BP positive stress granules in ASAP expressing cells within the same time course of light up to 30 minutes (Extended Data Fig. 6r-u).

**Figure 6:**
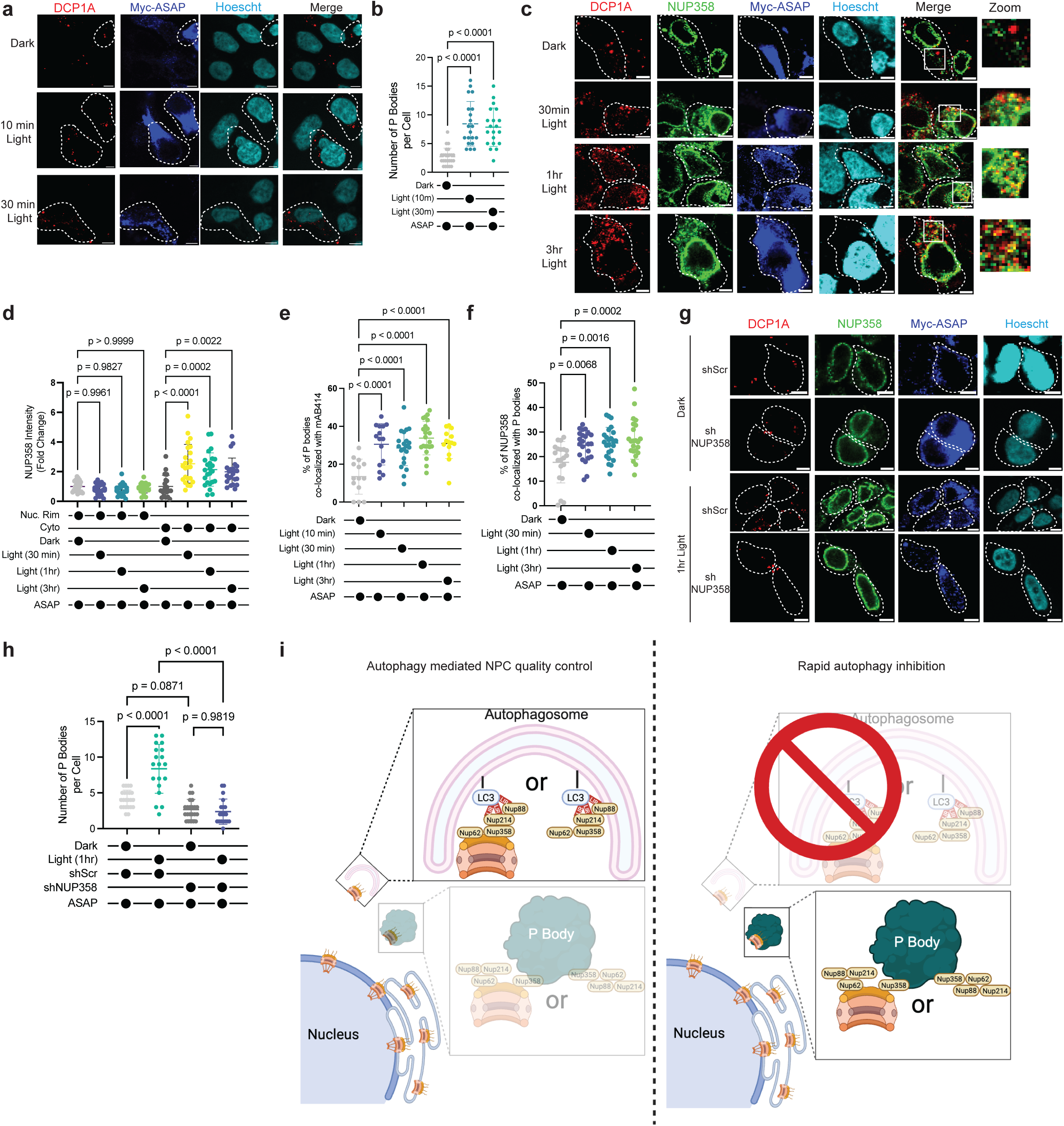
Nucleoporins accumulate in P bodies after autophagy inhibition via NUP358. **(a-b)** HEK293T cells transfected with ASAP and treated with 450nm pulsed light for indicated periods of time followed by IF. **(a)** Representative IF images of the P body marker, DCP1A, and Myc-tag for ASAP. Scale bar = 5μM. **(b)** Quantification of the number of P bodies per cell. N=21 cells per condition. Data are graphed as mean ± S.D. and are representative of 3 independent experiments. Statistical analyses were performed using a One Way ANOVA. **(c-d)** HEK293T cells transfected with ASAP and treated with 450nm pulsed light for indicated periods of time followed by IF. **(c)** Representative IF images for DCP1A, NUP358, and Myc-tag for ASAP. Scale bar = 5μM. **(d)** Quantification of the NUP358 signal that is cytoplasmic or on the nuclear rim, data graphed as fold change to the respective dark condition for cytoplasmic and rim signal. N=19-33 cells per condition. Data are graphed as mean ± S.D. and are representative of 2 independent experiments. Statistical analyses were performed using a One Way ANOVA. **(e)** HEK293T cells transfected with ASAP and treated with 450nm pulsed light for indicated periods of time followed by IF. Quantification of the percent of P bodies that co-localize with cytoplasmic MAb414 signal. N=21-33 cells per condition. Data are graphed as mean ± S.D. and are representative of 2 independent experiments. Representative IF images DCP1A, MAb414, and Myc-tag for ASAP shown in Extended Data Fig. 7a. Statistical analyses were performed using a One Way ANOVA. **(f)** HEK293T cells transfected with ASAP and treated with 450nm pulsed light for indicated periods of time followed by IF. Quantification in the percent of cytoplasmic NUP358 signal that colocalizes with DCP1A P body signal. N=19-33 cells per condition. Data are graphed as mean ± S.D. and are representative of 2 independent experiments. Statistical analyses were performed using a One Way ANOVA. Representative images are shown in in Fig. 6c. (**g-h**) HEK293T cells with stable expression of shScramble (shScr) or shNUP358 and transfected with ASAP and treated with 450nm pulsed light for indicated periods of time followed by IF. **(g)** Representative IF images for DCP1A, NUP358, and Myc-tag for ASAP. Scale bar = 5μM. **(h)** Quantification of the number of P bodies per cell. N=19-23 cells per condition. Data are graphed as mean ± S.D. and are representative of 3 independent experiments. Statistical analyses were performed using a One Way ANOVA. **(i)** Graphical abstract of the model showing autophagy degrades cytoplasmic nucleoporins under basal conditions when autophagy is blocked, and undegraded nucleoporins aggregate with P bodies. This autophagic degradation could target cytoplasmic aggregates of nucleoporins or annulate lamellae.

NUP358 (also known as RanBP2), one of the NPC proteins that co-precipitates with LC3 (Fig. 5), was previously shown to be important for P body formation and interact with proteins found in P bodies^57, 64^. IF with antibodies specific for NUP358 shows that it also accumulates in the cytoplasm immediately after light mediated autophagy inhibition (Fig. 6c-d) Given that rapid autophagy inhibition causes an increase in both P bodies and an accumulation of nucleoporins, co-IF was performed to determine interactions between the two. Indeed, just 10 minutes of light increases the percentage of P bodies that colocalize with nucleoporins, specifically NUP358 (Fig. 6c,e, Extended Data Fig. 7a-d). Likewise, the percentage of MAb414 signal, and especially NUP358, that colocalizes with P bodies is increased in ASAP expressing cells (Fig. 6f, Extended Data Fig. 7e). P bodies specifically colocalize with nucleoporins and not indiscriminately with the ER (Extended Data Fig. 7f-g). To test if NUP358 is required for light-induced P body accumulation in ASAP expressing cells, we silenced NUP358. Since NUP358 is an essential gene^65^, only partial knock down (∼50%) is tolerable in these cells (Extended Data Fig. 7h). This level of knock down results in a measurable decrease in cytoplasmic NUP358 via IF, although NUP358 signal on the nuclear rim is not affected, and prevents light-induced accumulation in cytoplasmic NUP358 (Fig. 6g, Extended Data Fig. 7i). Silencing NUP358 also prevents light-induced P body accumulation (Fig. 6g-h), indicating that NUP358 is required for, and upstream of, autophagy inhibition-induced P body accumulation.

Together, our studies leverage ASAP to rapidly inhibit autophagy, revealing that autophagy constitutively degrades cytoplasmic complexes of nucleoporins via the LIRs in NUP214 and NUP88. Rapid autophagy inhibition leads to the accumulation of cytoplasmic nucleoporins including those that interact with NUP214 and NUP88 like NUP358. Cytoplasmic accumulation of NUP358 induces P body accumulation as an additional stress response of immediate autophagy inhibition. Therefore, when autophagy is inactive, cytoplasmic nucleoporins instead accumulate in these induced P bodies, via interaction with NUP358 (Fig. 6i).

## LIMITATIONS

ASAP is a powerful tool for rapid manipulation of autophagy; however, there are some limitations. ASAP requires transiently transfected plasmids which prohibits the use of ASAP for long term experimentation. There are also several cell culture systems that are not amenable to transient transfection that limit the use of ASAP. Most optogenetics systems described have some leak in the system. While we confirmed very little off-target TEV cleavage in the dark, it’s important to note that high expression of ASAP, irrespective of light, results in some LC3-II accumulation. This can be observed in some experiments here, although treatment with light always further increases LC3-II; nonetheless, high expression can prevent the tight regulation of the system.

## DISCUSSION

Together, these studies describe a novel optogenetic tool, ASAP, to rapidly block autophagy. There have been many advances in the field to measure autophagy across different systems, as well as unique tools to induce specific forms of autophagy^35, 66^. However, until now, autophagy blockade has been restricted to slow genetic methods or non-specific pharmacological inhibitors. ASAP represents a significant advance in the field, enabling acute, reversible autophagy inhibition with robust temporal precision. With a plasmid-based system, the tool can be expressed and applied across different models (Fig. 1, Extended Data Figs. 1, 2, 8) with potential for even more advanced spatial resolution via tissue-specific promoters. Furthermore, ASAP can be adapted for *in vivo* applications across various model organisms to investigate autophagy’s role in both healthy conditions, such as development, and diseased states, including neurodegeneration or cancer.

We benchmarked ASAP against pharmacological inhibition and show that while Baf-A1 takes 1-4 hours to inhibit autophagy, while ASAP blocks autophagy within 5 minutes of light exposure (Fig. 1, 2, Extended Data Figs. 1, 2). We also found that ASAP is reversible within 30 minutes of removal of the light, while autophagy levels do not revert to steady-state even 24 hours after Baf-A1 is washed out (Fig. 2). Moreover, given the genetic nature of ASAP, this tool is likely more specific than pharmacological autophagy inhibition, although it is important to note that STX17 has been shown to localize to mitochondria^67, 68^, so there could be additional on-target side effects in ASAP expressing cells. Prior studies have used doxycycline inducible systems to inhibit autophagy^30, 69^ or CRISPR/Cas9 to knock out autophagy genes^70^, both of which take several days to take effect. Therefore, we conclude that ASAP is faster and more reversible than prior tools used to inhibit autophagy. While it is difficult to directly compare the efficiency of ASAP to other systems, the data here show that minutes of light leads to the same effects, on both autophagy inhibition and substrate accumulation, seen with hours of Baf-A1 suggesting that ASAP can block autophagy to the same efficiency, but faster.

With the introduction of this new tool in cells, we combined ASAP with quantitative proteomics to identify the most robust autophagy substrates, such as proteins that begin accumulating within minutes of autophagy inhibition (Fig. 3, Extended Data Fig. 3). While we identified several known autophagy cargo adapters and substrates, we also discovered many proteins previously not recognized as autophagy substrates in mammalian cells. Importantly, we acknowledge that proteomics is limited by peptide abundance, and thus our analyses may be missing low-abundance proteins. Nonetheless, proteomics, followed by confirmatory immunoblotting, reveal that mitochondrial proteins, nuclear pore complex proteins, and select protein condensates increase immediately after autophagy inhibition.

Of these top hits, the nuclear pore complex and its relationship to protein condensates had not previously been linked to autophagy in mammalian systems. However, NPC-autophagy was previously described in Saccharomyces cerevisiae where NUP159 plays an important role in mediating interactions with LC3 through a LIR sequence to facilitate nitrogen starvation-induced degradation of the nuclear pore complex via autophagy^50–52^. NUP159 is an ortholog of the human NUP214 NPC protein and we demonstrate here that NUP214, along with NUP88 have evolutionarily conserved and bona fide LC3-interacting regions as well (Fig. 5, Extended Data Fig. 5). Therefore, our studies indicate that nucleoporinphagy, first described in yeast, is conserved in mammalian cells. The data shows that several nucleoporins immunoprecipitate with NUP214 and NUP88, suggesting that multiple cytoplasmic nucleoporins are in complex together and therefore also get degraded by autophagy. Interestingly, many of the nucleoporins found in ER localized NPCs, known as annulate lamellae (AL), appear to interact with LC3 and accumulate more in the cytoplasm after autophagy inhibition, while NUP153, which is excluded from AL, had a more muted phenotype in the cytoplasm (Figs 4, 5, Extended Data Figs. 4,5). Together, these results suggest that cytoplasmic nucleoporins found in AL are potentially targeted by autophagy more than other cytoplasmic nucleoporins and more than nucleoporins on the nuclear envelope that are part of functional NPCs. However, electron microscopy is required to determine if autophagy targets intact AL or simply nucleoporin aggregates that are destined for AL.

Our studies reveal that nucleoporinphagy is constitutively active under basal conditions, likely to maintain quality control of newly synthesized nucleoporins prior to their incorporation into functional NPCs embedded in the nuclear envelope. This is supported by immunofluorescence imaging showing that immediately after autophagy inhibition, cytoplasmic nucleoporins accumulate more than those already embedded in the nuclear envelope, and the cytoplasmic accumulation is blocked in the presence of CHX (Fig. 4, Extended Data Fig. 4). The sorting mechanisms that determine which nucleoporins are incorporated into functional NPCs on the nuclear envelope, which form annulate lamellae, and which remain as pre-assembled aggregates in the cytoplasm are largely unknown. An important unanswered question from the current study is how autophagy preferentially targets the cytoplasmic nucleoporins instead of intact NPCs. One theory, yet to be tested, is that the LIRs may become less accessible when NUP214 and NUP88 are incorporated into NPCs protecting them from autophagic degradation.

These studies used proliferative immortalized HEK293T cells as a mammalian model. Therefore, this tight regulation of nucleoporinphagy could be especially prevalent in rapidly dividing cells that must maintain large pools of NPC reserves. While the data here indicate some changes in cell viability with ASAP (Fig. 2), future studies will need to characterize these mechanisms in cells with uncontrolled proliferative capacity, like cancer cells, and in non-dividing cells, like neurons.

The proteomics studies also suggested an increase in protein condensates, and immunofluorescence imaging revealed that P bodies, but not stress granules, are induced within minutes of autophagy inhibition (Fig. 3, 6, Extended Data Fig. 6). Given that many proteins known to be associated with P bodies do not have predicted LIR motifs, we postulate that P bodies are an immediate stress response to autophagy inhibition rather than an autophagy substrate. Our data show this phenomenon is mediated by NUP358 which accumulates in cytoplasmic NPCs after autophagy inhibition and is critical for P body formation^57, 64^ (Fig. 6, Extended Data Fig. 7). While the work of Sahoo et al^64^ suggests that P bodies could interact with cytoplasmic nucleoporins via the NUP358 SUMOylation motif, future studies are needed to fully elucidate the mechanism of interaction in the context of rapid autophagy inhibition. Beyond DCP1A and NUP358, P bodies also contain translationally repressed RNAs, suggesting that a rapid response to autophagy inhibition could lead to altered protein translation. Future studies are needed to determine if global protein translation is stalled after autophagy inhibition, potentially due to decreased amino acid pools, or if select RNA transcripts are associated with autophagy-inhibition-induced P bodies. The proteomics analyses also show many proteins that decrease immediately after light which should be further explored to determine if these changes could be regulated by altered protein translation and P bodies.

Together these studies describe a tool to target autophagy that can be broadly applied across different physiological contexts. Beyond identifying autophagy substrates, the proteomics data set from ASAP-expressing cells can be further explored to assess the immediate cellular response to autophagy inhibition, including both upregulated and down regulated proteins. Given the central role of autophagy in many different diseases, including neurodegeneration, cancer, and other aging-related conditions, these studies provide a much-needed advance that will help propel the field further toward targeting autophagy and its downstream effects to be treat disease.

## Acknowledgements

This work was supported by an NIH/NCI grant DP2 CA290705 (C.G.T), a Pew-Stewart Scholar award 00036071 (C.G.T), a CZI Leadership Award (C.G.T), a T32 CA009370 fellowship (L.R.P.), the HMMI Hanna H. Gray fellowship GT16776 (L.R.P.), A T32GM145427 fellowship (A.K.), The Salk Pioneer Fund postdoctoral scholar award (P.M.), Salk Women and Science Award 2021 (PM), 2023 (AK) and 2023 (PM), and the UC San Diego Health Sciences Collaborative Proteomics Center (D.J.G and I.W.). We also acknowledge the Waitt Advanced Biophotonics Core Facility of the Salk Institute (RRID:SCR_014838) with funding from NIH-NCI CCSG P30 CA014195, NIH-NIA San Diego Nathan Shock Center P30 AG068635, and the Waitt Foundation.

## Declaration of Interests

The authors declare no competing interests.

## Methods

### Molecular cloning of ASAP construct and light treatments

The fully annotated sequences for all ASAP and CTRL constructs are provided in Extended Data Fig. 8a-b. The ASAP plasmid was created using an N-terminal IgK-leader sequence adapted from pDisplay-mgSrtA which was a gift from P. Chen (Addgene plasmid # 125792; http://n2t.net/addgene:125792; RRID:Addgene_125792)^71^ and PCR amplifying the TM domain from a gene block synthesized by IDT. hLOV1 sequence was gifted by Jin Zhang lab at UCSD. STX17ΔNTD was cloned from pMRXIP-GFP-STX17ΔNTD which was a gift from Noboru Mizushima (Addgene plasmid # 89941; http://n2t.net/addgene:89941; RRID:Addgene_89941)^30^. p2A sequence was cloned using a gene block synthesized by IDT. TEV sequence was cloned from NLS-mCherry-LEXY (pDN122) which was a gift from Barbara Di Ventura & Roland Eils (Addgene plasmid # 72655; http://n2t.net/addgene:72655; RRID:Addgene_72655)^72^. The recombinant DNA was ligated into a N1 vector backbone using NheI and NotI restriction sites. The CTRL plasmid was created by PCR amplifying TM and eLOV and ligating into N1 vector using SalI and NotI restriction sites. ASAPΔtevS was created by removing tevS from the original ASAP construct, and ASAPΔTEV was generated by deleting the TEV protease sequence from the ASAP construct. The ASAP-mRuby2 construct was engineered by inserting the mRuby2 sequence (from the mRuby2-N1^73^ vector which was a gift from Michael Davidson, Addgene plasmid #54614; https://n2t.net/addgene:54614; RRID:Addgene 54614) between the TM and hLOV1 domains of the ASAP construct synthesized by Epoch Life Science. Similarly, the control-mRuby2 construct was created by inserting the mRuby2 sequence between the TM and eLOV domains in the original control construct. ASAP and CTRL were transfected into 500,000 - 800,000 cells in 35-mm dishes. Cells were 50-70% confluent on the day of transfection. HEK293T cells were transfected with 1 ug DNA and HCT116, NCI-H292 cells were transfected with 2.5 ug DNA due to their lower baseline transfection efficiencies. All dishes were wrapped in tinfoil to block ambient light while in culture. Next day, the dishes were placed in a light box inside the tissue culture incubator. The light box consisted of a 3.00″ L x 17.00″ W x 17.00″ D size box (BUD industries, Model # AC-1430) where the bottom was open and all sides were covered with LED light strips (Amazon ASIN B086TYT7MT) equipped to emit 450 – 480 nM light (Extended Data Fig. 8g-j). The LED lights were connected to a timer (Amazon ASIN B07D3BFY6R which was set to emit 0.6-0.7 mW/cm2 pulsed light at 10s on, 5s off intervals. To reduce ambient light exposure, ASAP transfected cells were handled in a closed tissue culture space lit by red light only. Cells were either harvested immediately after light treatment for downstream assays or wrapped in tinfoil and placed back in the incubator. Once cells were lysed, all downstream assays were performed on the normal lab bench under ambient light conditions.

### Cell Culture and transfections

All cell lines were maintained at 37°C and 5% CO2. HCT116, HEK293T, and NCI-H292 cell lines were obtained from the laboratory of Dr. Andrew Thorburn at the University of Colorado. HEK293T and HCT116 cells were cultured in Dulbecco’s Modified Eagle’s Medium (DMEM, Thermo Fisher Scientific) with 10% fetal bovine serum (FBS, Omega Scientific Inc.) and 1% Penicillin Streptomycin (Pen/Strep; Thermo Fisher Scientific). NCI-H292 Cells were cultured in Roswell Park Memorial Institute medium (RPMI 1640; Thermo Fisher Scientific) with 10% FBS (Omega Scientific) and 1% Pen/Strep (Thermo Fisher Scientific). All cell lines were periodically monitored for mycoplasma contamination with the MycoAlert Mycoplasma Detection Kit (Lonza). Cell transfections were performed using Opti-MEM and Lipofectamine 3000 (Thermo Fisher Scientific, Cat# L300015) along with plasmid DNA in 35mm tissue culture treated dishes. Cells were harvested for downstream assays the next day. Drug treatments include Bafilomycin-A1 (10nM, Sigma, catalog #B1793), MG132 (25 uM, Selleckchem, catalog #S2619), Puromycin (100μg/mL, Thermo fisher scientific, catalog #A1113803), Sodium Arsenite (0.5 mM, Santa Cruz Biotechnology sc-301816), Cycloheximide (100ug/ml; Sigma Aldrich; cat#C4859-1ML). To knock down NUP358 lentivirus were generated using the pPLKO.1 containing shRNA sequences ATGTACTCCTGACGTATTAAA for RanBP2 and CCGGGCGCGATAGCGCTAATAATTTCTCGAGAAATTATTAGCGCTATCGCGCTTTTT for a scrambled NT control from Millipore Sigma, and added to HEK293T cells followed by puromycin selection to create pooled populations of cells with stable knock down.

### Immunofluorescence imaging

Coverslips (12mm, Fisher Scientific NC1117320) were placed into 35mm dishes and treated with UV light for 20 minutes, then incubated with Poly-L-Lysine (Cell Biologics, Cat#NC1468070) for 1 hour followed by three PBS washes. 500,000 cells were seeded per plate. The next day, cells were transfected with 1 ug desired plasmids. Next day, after the light exposure or drug treatment, the media was removed, followed by a 15-minute incubation in Formalin (Cat #SF100) and washed again 3X with PBS followed by permeabilized in 0.1% Triton-X in PBS for five minutes followed by 3X washes with PBS. For IF experiments with NPC proteins, cells were fixed in 4% PFA for 5 minutes followed by 3X washes and incubationin blocking buffer for 1 hour. For NPC proteins the blocking buffer included 1x PBS with 10mg/ml BSA, 0.1% Triton X-100, and 0.02% SDS. For all other antibodies the blocking buffer included 5% normal goat serum (NSG)/0.3 M glycine/PBS. Cells were then washed 3X with PBS and incubated in the primary antibody diluted in blocking buffer overnight. Primary antibody dilutions include LC3 (1:2500, Cells Signaling Technology Cat#3868), SERCA2 (1:1000, Thermo Scientific, Cat#MA3-919), NUP88 (1:1000, BD Biosciences, Cat#611896), NUP214 (1:1000, Abcam, Cat#ab70497), MAb414 (1:1000, BioLedgend Cat#902902), NUP93 (1:1,000; Santa Cruz Biotechnology; Cat# sc-374399), NUP153 (1:1,000; Abcam; Cat# AB84872), Myc (1:1000, Thermo Scientific, Cat# A-21281), DCP1A (1:2000, Thermo Fisher Scientific, Cat#22373-1-AP), DCP1A (1:1000, Abnova cat#H00055802-M06) and G3BP (1:500, Thermo Fisher Scientific, Cat#66486-1-IG). Cells were washed 3X with PBS, and the fluorescent secondary antibody was added and incubated for one hour. Secondary antibodies include goat anti-rabbit (1:1000, Thermo Scientific, Cat# A11008 or Cat# A11012), goat anti-mouse (1:1000, Thermo Scientific, Cat# A28175 or Cat# A11032), goat anti-chicken (1:1000, Thermo Scientific, Cat# A-21449). To label the plasma membrane, Wheat Germ Agglutinin conjugated to Alexa Fluor 488 was used (Thermo Fisher Cat# W11261). Following 3X washes with PBS, cells were incubated in Hoechst (diluted 1:2,0000 in PBS) for 15 minutes. Cells were washed 3X with PBS and then the coverslip was mounted onto a microscope slide Cat #22-037-246) using Vectashield mounting media (Cat # H-1700). Slides were imaged on an Olympus FV3000 confocal microscope and Z-stacks obtained. For live-cell imaging with mRuby2-ASAP, HEK293T cells were plated on glass bottom plates and transfected with mRuby2-ASAP were stained with ER-Tracker Green (Thermo Fisher Cat# E34251) for 30 min prior to live cell imaging with Olympus FV3000 confocal microscope using the PL APO 60X TIRF OIL objective NA1.50. *Image Quantification:* For SERCA2 and Myc co-localization, the JaCoP ImageJ plugin was used. To measure signal intensity for nucleoporins, a z plane with a clear nuclear rim was used and within Image J region of interests that include or exclude the nucleus were used to quantify signal intensity on the nuclear rim versus the cytoplasm. ImageJ was used to identify P body and stress granule puncta. After max-projection of Z-stacked images and thresholding, Particle Analyzer was used to count puncta (size: 5 – infinity pixels). To quantify co-localization of P bodies and MAb414 or LC3, Image Calculator with “AND” function within ImageJ was used. At least 15 cells were quantified per condition in every independent experiment, but if more cells were imaged per field of view, additional cells were included. To quantify mitochondrial phenotypes including count and branch length, Mitochondria Analyzer Plugin on ImageJ was used.

### Autophagy flux using mCherry-GFP-LC3

HCT116 cells with stable expression of pBabe-mCherry-GFP-LC3 (kindly gifted to us from Dr. Jay Debnath) were plated on glass bottom dishes (Mattek Cat # P35GC-1.5-14-C). Cells were transfected with ASAP or CTRL and treated with indicated periods of pulsed light using the light box. After light treatment, cells were either fixed with formalin for 15 min followed by immunofluorescence or prepared for live cell imaging using the Olympus FV3000 and Z-stacks were collected. Images were quantified using ImageJ. After max-projection of Z-stacks, signal in the red and green channels was thresholded and overlapping signal determined using the Image Calculator “AND” operation. Yellow puncta were quantified with Particle Analyzer (size: 0.02 – infinity and circularity: 0.1 – 1.0).

### FRAP

For nuclear transport measurements, mRuby2 fluorescence protein was inserted into ASAP and CTRL plasmid after TM transmembrane domain. ASAP-mRuby2 and CTRL-mRuby2 plasmids in N1 vector were constructed by Epoch Life Science company. 50,000 HEK293T cells stably expressing the transport reporter NLS-2xGFP-NES were plated in polymer coverslip 8-well µ-Slide dishes (ibidi Cat# 80824), as described previously^58^. Cells were transiently transfected with 100ng ASAP-mRuby2 or CTRL-mRuby2 and exposed to 450 nm pulsed light for 3 h prior to imaging. For FRAP experiments with 48-72 h pulsed light treatment, 30,000 cells were transfected with 50 ng DNA. Cells were imaged using a 40x objective on a Leica DMi8 microscope with a 37C and 5% CO2 stage top incubator (H301, Okolab) controlled by the Oko-Touch (Okolab). For FRAP, single z-stack images were acquired throughout the experiment. Three pre-bleach images were taken at 520 ms intervals to establish baseline fluorescence, followed by photobleaching of the nuclear region using 90% laser intensity for six consecutive iterations (520 ms per iteration). Fluorescence recovery was monitored for 180 s with images acquired every 3s using the Leica LAS X software. A total of 15–21 cells were imaged per condition and data was analyzed with ImageJ. Nuclear import data were calculated by dividing the nuclear and cytoplasmic background-subtracted signals and then normalized to the pre-bleach ratio.

### Molecular Cloning

Mutations in the LIR domain of NUP214 and NUP88 were made using Q5 site-directed mutagenesis kit (New England Biolabs, E0554S) following manufacture’s protocol. The NUP214 double mutant construct was designed through Epoch Life Sciences. The primers for point mutagenesis were designed using NEBasechanger website (https://nebasechanger.neb.com).

### Western blotting (densitometry quantification, antibodies used and dilutions)

Whole cell lysate samples were washed with cold PBS and harvested on ice with RIPA buffer (4M NaCl, 5ml IGEPAL, 17 mM Sodium Deoxycholate, 10% SDS, 1M tris ph7.0, 0.5 M NaF, 0.5M EDTA pH 7.0) and protease inhibitor cocktail (Roche, Cat #11873580001). Lysates were sonicated and spun down at 20,000xg for 15min at 4°C, after which membrane pellet was discarded. Protein concentration was measured from the supernatant and determined via Bradford assay. 3X Laemmli Sample buffer (4X Laemmli sample buffer, Bio-Rad, Cat#1610747) and 1/10 β-mercaptoethanol (Sigma, Cat# M625) was added and samples boiled at 95°C. Western blotting was performed using standard methods including protein separation on SDS-PAGE 1.5mm mini gels in running buffer (250 mM tris base, 190 mM glycine, 3.4 mM SDS) at 100V for 2hrs, followed by transfer to PVDF membranes in transfer buffer (25 mM tris base, 180 mM glycine) using a semi-dry transfer apparatus at 15V for 70 minutes. Membranes were blocked in 5% milk in TBST (TBST: 20 mM tris acid, 1.3 mM tris base, 100 mM NaCl, and Tween20) for 1hr at room temperature with gentle rocking, washed twice in TBST, and then incubated overnight at 4°C with gentle rocking in primary antibodies. Antibodies include LC3 (1:2500, Novus Biologics, Cat# NB100-2220SS), SQSTM1/p62 (1:5000, Novus Biologics, Cat# H00008878-M01), Myc (1:2000, from Colorado), MAb414 (1:1000, Biolegend, Cat# 902902), NUP214 (1:1000, Proteintech, Cat# 24113-1-AP), NUP88 (1:1000, Proteintech, Cat# 55465-1-AP), TOMM20 (1:1000, Cell Signaling Technology (CST), Cat# 42406), TOM70 (1:1000, CST, Cat# 65619), BNIP3 (1:1000, CST, Cat# 44060), NIX (1:1000, CST, Cat# 12396), FUNDC1 (1:1000, CST, Cat# 49240), PDH (1:1000, CST, Cat# 3205), TIMM50 (1:1000, Abcam, Cat# AB23938), Tax1BP1 (1:1000, CST, Cat# 5105), OPTN (1:1000, CST, Cat# 70928), NDP52 (1:1000, CST, Cat# 60732), β-Actin (1:10,000, CST, Cat# 3700). Membranes were then washed three times in TBST and incubated for 1hr at room temperature with gently rocking in secondary antibodies followed by 3 more TBST washes. Secondary antibodies include Anti-Mouse IgG HRP-linked (CST 7076S), Anti-Rabbit IgG HRP-linked (CST 70742). Membranes were developed with Immobilon Western chemiluminescent HRP substrate (Sigma-Aldrich, WBKLS0500) and analyzed on the ChemiDoc imaging system (Bio-Rad). Band intensity was calculated using ImageJ and expressed as a ratio between protein of interest to loading control. Experimental samples were then normalized to controls

### Co-Immunoprecipitation

4 x 10^6^ cells were seeded in a 10cm plates. Whole cell lysates were harvested in 0.5 mL of 1X cell lysis buffer (CST, Cat#9803) after being transfected with 8 μg of the desired plasmids as described above. For co-IPs, pCMV 3xFLAG-LC3B WT was a gift from Robin Ketteler (Addgene plasmid # 123092; http://n2t.net/addgene:123092; RRID:Addgene_123092)^74^, NUP88 (Origene SKU SC310134) or NUP214 (pcDNA-DEST53-GFP-NUP214; plasmid made by Martin Hetzer lab). The corresponding empty vector (EV) controls were also used. Of the 0.5 mL of lysate harvested, 200 μL of lysate was incubated with antibody for IP and 200 μL was incubated with the corresponding IgG (Rabbit IgG CST Cat#2729, Mouse IgG CST Cat#53484) at 4°C overnight with mixing. The remaining 100 μL was kept at -20°C for western blot. Protein A magnetic beads (CST Cat#73778) for rabbit pull down and Protein G magnetic beads (CST Cat#70024) for mouse IgG pull down was used. The lysate was pre-cleared by incubating it with beads without antibody for 1hr. Magnetic beads were washed and then incubated with the IgG and target antibody lysates for one hour at RT. Then, a magnetic stand was used to collect the flow from the tubes and saved for western blot analysis. Each tube was washed 3-5 times with 1X cell lysis buffer. 3X SDS sample buffer was prepared fresh by adding 1/10 volume 30X DTT (1.25 M DTT, CST#14265) to 1 volume of 3X SDS loading buffer (CST Cat#7723). 50ul of the sample buffer was added to the washed beads and incubated at 96-100°C for 5 minutes. 5-20 ul elute was used for western blotting as described above.

### Cell fractionation

Cells plated in a 10cm dish were transfected with ASAP and either kept in the dark or exposed to pulsed blue light for 1h were washed with PBS, subjected to trypsin, and collected by centrifugation at 300g x 5min. Cell pellets were resuspended in 2ml of RSB Hypotonic Buffer (10 mM NaCl, 1.5 mM MgCl2, 10 mM Tris-HCl (pH 7.5)) for 10 minutes to induce cell swelling. Subsequently, the cell suspension was transferred to a 5ml Potter-Elvehjem homogenizer and subjected to 20 strokes with the pestle, making sure to avoid forming bubbles. Afterwards, 1ml of deionized H2O and 2ml of 2.5X MS Homogenization buffer (525 mM mannitol, 175 mM sucrose, 12.5 mM Tris-HCl (pH 7.5), 2.5 mM EDTA (pH 7.5)) were added to the homogenizer, the top of the homogenizer was sealed with paraffin, and the homogenizer was inverted 10 times to mix the cell lysate. 500ul of the solution was collected as a whole cell lysate (WCL) while the rest of solution was split into 1.5ml microcentrifuge tubes. The collected solution was spun at 1300g x 5min. The subsequent pellet was resuspended in RIPA buffer and defined as an organelle-enriched fraction. The supernatant from this spin was further centrifuged at 7000g x 15min. Following this spin, the pellet was discarded, and the supernatant was collected as a total cytosol fraction. The WCL, organelle-enriched fraction, and the cytosol fraction were flash-frozen, sonicated, and spun down at 21000g x 10min. The protein concentration of each sample was quantified via Bradford assay and 10ug of each sample was loaded on a 12% SDS-PAGE assay with a subsequent western blot performed to evaluate the indicated proteins.

### Incucyte live cell imaging

Live cell imaging was performed with the Incucyte S3 (Sartorius) at 10X magnification. HCT116 cells containing nucleored were seeded in 96-well plates. The next day CellROx (5 μM) or Cell Event Caspase3/7 (5 μM) was added. Images were taken every 4hrs for CellROx. For CellEvent™ Caspase-3/7 experiments, plates were removed from the Incucyte and exposed to pulsed light for 30 min in an optogenetic incubator, then returned to the Incucyte for imaging 10 min after light exposure. After an additional 20 min, plates were removed for the next 30 min light-pulse cycle. This cycle was repeated for a total duration of 10hrs. Cell count, Cell Rox, and Caspase3/7 were measured using Incucyte software. For cell viability experiments in ASAP expressing cells, transfected cells were exposed to pulsed blue light for 1hr for short term time points, followed by pulsed light for 5 minutes every 30 minutes for 3 days for longer time points. One-time scans were taken every 24hrs in the Incucyte.

### Mass Spectrometry for TMT proteomics

#### Sample Preparation for Mass Spectrometry Analysis

Sample denaturation, reduction, and alkylation: Samples were suspended in 500 µL of Lysis buffer composed of 6M urea [Fisher Chemicals, CAT# U15-500], 7% SDS [Fisher BioReagents, CAT# BP166-500], 50 mM triethylammonium bicarbonate (TEAB) [Sigma-Aldrich, CAT# T7408-500ML], pH 7.7; supplemented with cOmplete™ ULTRA Tablets, EDTA-free (1 tablet for 20 mL of buffer) [Roche, CAT# 05892791001] and PhosSTOP™ (1 tablet for 20 mL of buffer) [Roche, CAT# 4906845001]. Samples were transferred into 2 mL Protein LoBind Tubes [Eppendorf, CAT# 022431102], incubated for 10 minutes in a Vortex-Genie 2 vortex mixer [Scientific Industries, CAT# 00-SI-0236], and sonicated twice using Qsonica Sonicator Q125 [Qsonica LLC, CAT# Q125-110] equipped with a 1.6 mm microtip probe [Qsonica LLC, CAT# 4417] at a following setting: amplitude 20; 1-second pulse; 5-second break; 5 pulses. Lysates were centrifuged for 5 minutes at 16,000×g, room temperature. Supernatants were transferred into new 2 mL Protein LoBind Tubes, and proteins were chemically reduced by the addition of 5 µL of 0.5 M dithiothreitol (DTT) [Invitrogen, CAT# 15508-013] and 30-minute incubation at 47. Next, samples were cooled on ice for 5 minutes, and proteins were alkylated by the addition of 15 µL of 0.5 M iodoacetamide [Sigma-Aldrich, CAT# I1149] and 45-minute incubation at room temperature in the dark. The alkylation reaction was stopped by adding 5 µL of 0.5 M DTT and 5 minutes of incubation at room temperature. Samples were transferred and stored at -80.

#### Protein digestion

Samples were thawed at room temperature and centrifuged for 5 minutes at 16,000×g, room temperature. Supernatants containing solubilized chemically reduced and alkylated proteins were transferred into new 2 mL Protein LoBind Tubes and acidified by the addition of 54 µL of 12% phosphoric acid [Sigma-Aldrich, 49685-500ML] and mixed with 1500 µL of Binding buffer composed of 90% methanol [Fisher Chemicals, CAT# A452-4], 50 mM TEAB, pH 7.1. Samples were next loaded, ∼700 µL at a time, onto S-Trap™ mini columns [ProtiFi, CAT# C02–mini-80] by 1-minute centrifugation at 4000×g, room temperature. Columns were washed 5 times with 500 µL of Binding buffer by 1-minute centrifugation at 4000×g, room temperature; centrifuged for 5 minutes at 4000×g, room temperature, to remove residual methanol; and transferred into new collection tubes. Trypsin digestion mix, composed of 20 µL (10 µg) of Sequencing Grade Modified Trypsin [Promega, CAT# V5113] and 105 µL of 50 mM TEAB, was loaded into each column by brief centrifugation (<2000×g) and reapplication of the resulting flow-through. Next, the proteins were on-column trypsin-digested for 3 hours at 47. Peptides were eluted with 125 µL of 50 mM TEAB (1-minute centrifugation at 2000×g, room temperature), 125 µL of 5% formic acid [Fisher Chemicals, CAT# A118P-500] (1-minute centrifugation at 4000×g, room temperature), and 125 µL of 50% acetonitrile [Fisher Chemicals, CAT# A998-4] (5-minute centrifugation at 4000×g, room temperature), and transferred into 2 mL Protein LoBind Tubes. Obtained peptides were next frozen at -80 and dried using Savant SPD111V SpeedVac Concentrator [Thermo Fisher Scientific, CAT# SPD111V-115], in line with Savant RVT5105 Refrigerated Vapor Trap [Thermo Fisher Scientific, CAT# RVT5105-115].

#### Peptide desalting

Dried peptides were suspended in 500 µL of 0.1% trifluoroacetic acid (TFA) [Thermo Fisher Scientific, CAT# 28901] and incubated for 20 minutes in a Vortex-Genie 2 vortex mixer. Sep-Pak tC18 1 cc Vac Cartridges (50 mg sorbent) [Waters, CAT# WAT054960] were placed in a NucleoVac 24 Vacuum Manifold [MACHEREY-NAGEL, CAT# 740299] and washed via laboratory vacuum with 1 mL of 100% acetonitrile and 2 mL of 0.1% TFA. Peptide solutions were centrifuged for 5 minutes at 16000×g, room temperature, and the supernatants were applied onto the tC18 cartridges. Column-bound peptides were washed with 5 mL of 0.1% TFA and eluted with 750 µL of 40% acetonitrile, 0.5% acetic acid; and 750 µL of 80% acetonitrile, 0.5% acetic acid into a 2 mL Protein LoBind Tubes. Desalted peptides were frozen at -80 and dried using Savant SPD111V SpeedVac Concentrator in line with Savant RVT5105 Refrigerated Vapor Trap.

Peptide quantification and TMT-labeling: Dried peptides were resuspended in 750 µL of 50% acetonitrile and incubated for 20 minutes in a Vortex-Genie 2 vortex mixer. Pierce Quantitative Colorimetric Peptide Assay kit [Thermo Fisher Scientific, CAT# 23275] was used according to the manufacturer’s recommendations to determine peptide concentration. “Bridging” samples, allowing for later data normalization between the TMTpro 16-plexes were prepared by combining 8 µg of each sample (total 240 µg per “Bridging” samples mix). Single 50 µg aliquots of each sample and four 50 µg aliquots of “Bridging” samples were transferred into 2 mL Protein LoBind Tubes, frozen at -80, and dried using Savant SPD111V SpeedVac Concentrator in line with Savant RVT5105 Refrigerated Vapor. Peptides were suspended in 50 µL of Resuspension buffer composed of 200 mM HEPES (BP310-500, Fisher BioReagents), 30% acetonitrile, and incubated for 20 minutes in a Vortex-Genie 2 vortex mixer. Two of “Bridging” samples were labeled with the TMTpro-126, and the remaining two with TMTpro-134 mass tag, while all other samples were randomized using the Microsoft Excel function “=SORTBY(name_range,RANDARRAY(COUNTA(name_range)))” for labeling with the remaining mass tags. Added 7 µL (20 µg) of TMTpro™ 16plex labels [Thermo Fisher Scientific, CAT# A44520, LOT# XE341501 and XL355490 (mass offsets for individual tags are provided in the quantification method file uploaded to MassIVE Repository)] to each sample and incubated for 1 hour at room temperature. The labeling reaction was terminated by adding 8 µL of 5% hydroxyl amine [Sigma-Aldrich, CAT# 438227-50ML] and a 15-minute incubation at room temperature. Samples were next acidified by adding 50 µL of 1% TFA. Samples labeled with each TMTpro™ 16plex set were combined in 2 mL Protein LoBind Tubes, frozen at -80, and dried using Savant SPD111V SpeedVac Concentrator in line with Savant RVT5105 Refrigerated Vapor.

High pH Reversed-Phase Peptide Fractionation: Dried samples were resuspended in 110 µL of 25 mM ammonium bicarbonate (ABC) [Spectrum Chemical, CAT# A1125], incubated for 10 minutes in a Vortex-Genie 2 vortex, and transferred to 300 µL Target Polyspring glass Inserts [Thermo Fisher Scientific, CAT# C4010-630] within 9 mm glass autosampler vials with pierceable PFTE septa [Thermo Fisher Scientific, CAT# C5000-580W]. Samples were fractionated by reverse-phase high pH liquid chromatography using UltiMate 3000 HPLC and UHPLC System with BioBasic™ 18 HPLC Column (4.6 mm x 250 mm) [Thermo Fisher Scientific, CAT# 72105-254630]. Solvent A of the mobile phase consisted of 5% acetonitrile and 10 mM ABC, while solvent B consisted of 90% acetonitrile and 10 mM ABC. Samples were separated into 96 fractions, in a 96-well format, by a 70.2-minute method composed of the isocratic flow of solvent A (2-minute duration), a linear gradient of 0-10% solvent B in solvent A (1-minute duration), a linear gradient of 10-40% solvent B in solvent A (57-minute duration), a linear gradient of 40-100% solvent B in solvent A (1-minute duration), an isocratic flow of 100% solvent B (4-minute duration), a linear gradient of 100-0% solvent B in solvent A (1-minute duration), and an isocratic flow of solvent A (4.2-minute duration), at a flow rate of 0.5 mL/min. Samples were collected starting from minute 11 till the end of the run. Fractions were combined within each column of the 96-well format to obtain a total of 12 fractions. A 1 mL aliquot of each concatenated fraction was transferred into 2 mL Protein LoBind Tubes, frozen at -80, and dried using Savant SPD111V SpeedVac Concentrator in line with Savant RVT5105 Refrigerated Vapor.

#### Sample reconstitution for mass-spectrometry analysis

Dried samples were suspended in 20 µL of 5% acetonitrile and 5% formic acid and mixed for 20 minutes in a Vortex-Genie 2 vortex. Samples were transferred to 300 µL Target Polyspring glass Inserts within 9 mm glass autosampler vials with pierceable PFTE septa.

#### Mass Spectrometry Data Acquisition

Proteomic spectral data was acquired with a Thermo Scientific Orbitrap Fusion Tribrid Mass Spectrometer. A 1 μL of each sample was separated by reverse-phase high pH liquid chromatography using a Thermo Easy-nLC 1200 liquid chromatography instrument (HPLC) [Thermo Fisher Scientific, CAT# LC140] and Easy-Spray PepMap Neo 2 µm C18 75 µm X 150 mm column [Thermo Fisher Scientific, CAT# ES75150PN]. Solvent A of the mobile phase consisted of 0.1% formic acid in HPLC-grade water, while solvent B consisted of 0.1% formic acid in acetonitrile. Samples were separated by a 180-minute method composed of a linear gradient of 6-25% solvent B in solvent A (duration 165 minutes), a linear gradient of 25-95% solvent B in solvent A (duration 5 minutes), and isocratic flow of 100% solvent B (duration 10 minutes), at a flow rate of 220 nL/min. MS acquisition was done in data-dependent mode at 60K resolution at a scan range of 500-1200 m/z for MS1. MS2 was collected using CID and MS3 using HCD.

#### Mass Spectrometry Data Search

Raw mass spectrometry files were searched using Proteome Discoverer 2.5.0.400. Precursor selection mass selection was set for 350-5000 Da. Data was searched against the UniProt reference proteome of Homo sapiens (Human) (accession UP000005640) with allowed 2 missed cleavages and a minimum peptide length of 6 amino acids and a maximum peptide length of 144. The false discovery rate was set to 5%. Precursor mass tolerance match was set to 10ppm and fragment mass tolerance of 0.6 Da. The “Bridging” samples were set in the “Sample type” category as “Control”. Normalization Mode was set to “Total peptide amount” and Scaling Mode was set to “On control averages”. Dynamic peptide modifications, natural and introduced during sample preparation, were added to the search: oxidation of methionine, acetylation of peptide N-terminus, loss of methionine at the peptide N-terminus, loss of methionine and acetylation at the peptide N-terminus, carbamidomethylation of cysteine, and TMTpro modification of peptide N-terminus and lysine residues.

#### Mass Spectrometry Data Statistical Analysis

Protein features with missing values were removed from further analysis. The average values were calculated for samples with technical replicates. Statistical significances (p < 0.05) between protein “Abundance (Scaled)” values were determined using Microsoft Excel’s built-in Student’s t-test with Welch’s Correction, where appropriate, based on an antecedent using Microsoft Excel’s built-in F test. Further stringency was imposed on binary comparisons by applying the multiple testing correction and calculating Benjamini-Hochberg adjusted p-values (q-values) using the R Studio ‘p.adjust()’ command.

### LIR prediction and sequence homology

iLIR autophagy database website (https://ilir.warwick.ac.uk) was used to identify predicted LIRs. Sequence homology was identified using BLAST function in Uniprot website (https://www.uniprot.org/blast).

### Quantitative Real Time PCR

RNA was harvested from samples using the Qiagen RNeasy Kit (Qiagen, Cat# 74134). cDNA was synthesized using 100 μg of RNA and the Bio-Rad iScript cDNA Synthesis Kit (Bio-Rad, Cat# 1708891). The qRT-PCR reaction was plated in Bio-Rad 96 well Hard-shell PCR plate (Bio-Rad, Cat# HSP9601), with a total volume of 20μL per well containing cDNA (diluted 1:50), forward and reverse primers and Fast SYBR Green Master Mix (Life Technologies, Cat# 4385610). The plate was sealed using the Bio-Rad microseal B (Bio-Rad, Cat# MSB1001) and spun down for 20 seconds. Plates were run on the Bio-Rad CFX Duet with the following cycles: 1 cycle at 95°C for 3 minutes, 40 cycles at 95°C for 10 seconds followed by 60°C for 30 seconds. A melt curve was performed at 65°C increased to 95°C in 0.5°C increments for 5 seconds each. All gene expression as normalized to a house keeping gene 18s. Primer sequences are described in Supplementary Table 2.

## Data Availability

All proteomics data (.raw file and data search information) are publicly available online at MassIVE Repository (https://massive.ucsd.edu) under identifier: MSV000099958, and at ProteomeXchange (https://www.proteomexchange.org) under identifier: PXD070944.

## Statistical Analyses

Statistical analyses were performed using GraphPad PRISM 10.0. When only two conditions were compared, an unpaired two-tailed T-test was performed. When more than two conditions were considered, a One-Way ANOVA was conducted followed by a Tukey’s multiple comparison test as indicated in the figure legends. All p values are reported within each figure. The Pearson’s correlation was calculated using JaCoP ImageJ plugin.

## Notes

### Competing Interest Statement

The authors have declared no competing interest.

### Summary of Updates

This version of the manuscript includes new experimentation included in almost all the figures and updated language in the text based on peer reviewer suggestions.

## References

1. Dikic, I. & Elazar, Z. Mechanism and medical implications of mammalian autophagy. Nat Rev Mol Cell Biol 19, 349–364 (2018).

2. Mizushima, N. The ATG conjugation systems in autophagy. Curr Opin Cell Biol 63, 1–10 (2020).

3. Yamamoto, H., Zhang, S. & Mizushima, N. Autophagy genes in biology and disease. Nat Rev Genet 24, 382–400 (2023).

4. Levine, B. & Kroemer, G. Biological Functions of Autophagy Genes: A Disease Perspective. Cell 176, 11–42 (2019).

5. Mizushima, N. & Levine, B. Autophagy in mammalian development and differentiation. Nat Cell Biol 12, 823–830 (2010).

6. Towers, C.G. & Thorburn, A. Therapeutic Targeting of Autophagy. EBioMedicine 14, 15–23 (2016).

7. Nah, J., Yuan, J. & Jung, Y.K. Autophagy in neurodegenerative diseases: from mechanism to therapeutic approach. Mol Cells 38, 381–389 (2015).

8. Sarkar, S., Davies, J.E., Huang, Z., Tunnacliffe, A. & Rubinsztein, D.C. Trehalose, a novel mTOR-independent autophagy enhancer, accelerates the clearance of mutant huntingtin and alpha-synuclein. J Biol Chem 282, 5641–5652 (2007).

9. Karsli-Uzunbas, G. et al. Autophagy is required for glucose homeostasis and lung tumor maintenance. Cancer Discov 4, 914–927 (2014).

10. Perera, R.M. et al. Transcriptional control of autophagy-lysosome function drives pancreatic cancer metabolism. Nature 524, 361–365 (2015).

11. Levy, J.M.M., Towers, C.G. & Thorburn, A. Targeting autophagy in cancer. Nat Rev Cancer 17, 528–542 (2017).

12. Assi, M. & Kimmelman, A.C. Impact of context-dependent autophagy states on tumor progression. Nat Cancer 4, 596–607 (2023).

13. Lopez-Otin, C., Blasco, M.A., Partridge, L., Serrano, M. & Kroemer, G. Hallmarks of aging: An expanding universe. Cell 186, 243–278 (2023).

14. Warr, M.R. et al. FOXO3A directs a protective autophagy program in haematopoietic stem cells. Nature 494, 323–327 (2013).

15. Ramirez-Pardo, I., et al. Chaperone-mediated autophagy sustains muscle stem cell regenerative functions but declines with age. Nat Metab (2025).

16. Mathew, R. et al. Functional role of autophagy-mediated proteome remodeling in cell survival signaling and innate immunity. Mol Cell 55, 916–930 (2014).

17. Anding, A.L. & Baehrecke, E.H. Cleaning House: Selective Autophagy of Organelles. Dev Cell 41, 10–22 (2017).

18. Kirkin, V., McEwan, D.G., Novak, I. & Dikic, I. A role for ubiquitin in selective autophagy. Mol Cell 34, 259–269 (2009).

19. Mancias, J.D., Wang, X., Gygi, S.P., Harper, J.W. & Kimmelman, A.C. Quantitative proteomics identifies NCOA4 as the cargo receptor mediating ferritinophagy. Nature 509, 105–109 (2014).

20. Sharma, K.B., et al. Quantitative Proteome Analysis of Atg5-Deficient Mouse Embryonic Fibroblasts Reveals the Range of the Autophagy-Modulated Basal Cellular Proteome. mSystems 4 (2019).

21. Zhou, X. et al. Integrated proteomics reveals autophagy landscape and an autophagy receptor controlling PKA-RI complex homeostasis in neurons. Nat Commun 15, 3113 (2024).

22. Russell, R.C. et al. ULK1 induces autophagy by phosphorylating Beclin-1 and activating VPS34 lipid kinase. Nat Cell Biol 15, 741–750 (2013).

23. Shpilka, T., Weidberg, H., Pietrokovski, S. & Elazar, Z. Atg8: an autophagy-related ubiquitin-like protein family. Genome Biol 12, 226 (2011).

24. Weidberg, H. et al. LC3 and GATE-16/GABARAP subfamilies are both essential yet act differently in autophagosome biogenesis. EMBO J 29, 1792–1802 (2010).

25. Kumar, S. et al. Mechanism of Stx17 recruitment to autophagosomes via IRGM and mammalian Atg8 proteins. J Cell Biol 217, 997–1013 (2018).

26. Itakura, E., Kishi-Itakura, C. & Mizushima, N. The hairpin-type tail-anchored SNARE syntaxin 17 targets to autophagosomes for fusion with endosomes/lysosomes. Cell 151, 1256–1269 (2012).

27. Hamasaki, M. et al. Autophagosomes form at ER-mitochondria contact sites. Nature 495, 389–393 (2013).

28. Takats, S. et al. Autophagosomal Syntaxin17-dependent lysosomal degradation maintains neuronal function in Drosophila. J Cell Biol 201, 531–539 (2013).

29. Tsuboyama, K. et al. The ATG conjugation systems are important for degradation of the inner autophagosomal membrane. Science 354, 1036–1041 (2016).

30. Uematsu, M., Nishimura, T., Sakamaki, Y., Yamamoto, H. & Mizushima, N. Accumulation of undegraded autophagosomes by expression of dominant-negative STX17 (syntaxin 17) mutants. Autophagy 13, 1452–1464 (2017).

31. Schrezenmeier, E. & Dorner, T. Mechanisms of action of hydroxychloroquine and chloroquine: implications for rheumatology. Nat Rev Rheumatol 16, 155–166 (2020).

32. Kim, M.W. et al. Time-gated detection of protein-protein interactions with transcriptional readout. Elife 6 (2017).

33. Golda, M. et al. P1’ specificity of the S219V/R203G mutant tobacco etch virus protease. Proteins 92, 1085–1096 (2024).

34. Marvin, J.S. et al. A genetically encoded fluorescent sensor for in vivo imaging of GABA. Nat Methods 16, 763–770 (2019).

35. Klionsky, D.J. et al. Guidelines for the use and interpretation of assays for monitoring autophagy (4th edition)(1). Autophagy 17, 1–382 (2021).

36. Towers, C.G., Wodetzki, D. & Thorburn, A. Autophagy and cancer: Modulation of cell death pathways and cancer cell adaptations. J Cell Biol 219 (2020).

37. Lee, S.Y. et al. Engineered allostery in light-regulated LOV-Turbo enables precise spatiotemporal control of proximity labeling in living cells. Nat Methods 20, 908–917 (2023).

38. Geng, L., Shen, J. & Wang, W. Circularly permuted AsLOV2 as an optogenetic module for engineering photoswitchable peptides. Chem Commun (Camb) 57, 8051–8054 (2021).

39. Bartolik, O. & Wang, W. A Single-Chain Light-Activatable Transcriptional Reporter for Fluorescently Tagging Mammalian Cells In Vitro. Chembiochem 27, e202500957 (2026).

40. Tsumagari, K., Isobe, Y., Imami, K. & Arita, M. Exploring protein lipidation by mass spectrometry-based proteomics. J Biochem 175, 225–233 (2024).

41. Pickles, S., Vigie, P. & Youle, R.J. Mitophagy and Quality Control Mechanisms in Mitochondrial Maintenance. Curr Biol 28, R170–R185 (2018).

42. Lazarou, M. et al. The ubiquitin kinase PINK1 recruits autophagy receptors to induce mitophagy. Nature 524, 309–314 (2015).

43. Narendra, D., Tanaka, A., Suen, D.F. & Youle, R.J. Parkin is recruited selectively to impaired mitochondria and promotes their autophagy. J Cell Biol 183, 795–803 (2008).

44. Padman, B.S. et al. LC3/GABARAPs drive ubiquitin-independent recruitment of Optineurin and NDP52 to amplify mitophagy. Nat Commun 10, 408 (2019).

45. Rossmann, M.P., Dubois, S.M., Agarwal, S. & Zon, L.I. Mitochondrial function in development and disease. Dis Model Mech 14 (2021).

46. Hurt, E. & Beck, M. Towards understanding nuclear pore complex architecture and dynamics in the age of integrative structural analysis. Curr Opin Cell Biol 34, 31–38 (2015).

47. Liu, J. & Hetzer, M.W. Nuclear pore complex maintenance and implications for age-related diseases. Trends Cell Biol 32, 216–227 (2022).

48. D’Angelo, M.A. & Hetzer, M.W. Structure, dynamics and function of nuclear pore complexes. Trends Cell Biol 18, 456–466 (2008).

49. Rabut, G., Doye, V. & Ellenberg, J. Mapping the dynamic organization of the nuclear pore complex inside single living cells. Nat Cell Biol 6, 1114–1121 (2004).

50. Allegretti, M. et al. In-cell architecture of the nuclear pore and snapshots of its turnover. Nature 586, 796–800 (2020).

51. Lee, C.W. et al. Selective autophagy degrades nuclear pore complexes. Nat Cell Biol 22, 159–166 (2020).

52. Tomioka, Y. et al. TORC1 inactivation stimulates autophagy of nucleoporin and nuclear pore complexes. J Cell Biol 219 (2020).

53. Hampoelz, B. et al. Nuclear Pores Assemble from Nucleoporin Condensates During Oogenesis. Cell 179, 671–686 e617 (2019).

54. Raghunayakula, S., Subramonian, D., Dasso, M., Kumar, R. & Zhang, X.D. Molecular Characterization and Functional Analysis of Annulate Lamellae Pore Complexes in Nuclear Transport in Mammalian Cells. PLoS One 10, e0144508 (2015).

55. Hampoelz, B. et al. Pre-assembled Nuclear Pores Insert into the Nuclear Envelope during Early Development. Cell 166, 664–678 (2016).

56. Lin, J. & Sumara, I. Cytoplasmic nucleoporin assemblage: the cellular artwork in physiology and disease. Nucleus 15, 2387534 (2024).

57. Lin, J. et al. RanBP2-dependent annulate lamellae drive nuclear pore assembly and nuclear expansion. Nat Commun 17 (2026).

58. Sakuma, S. et al. Homeostatic regulation of nucleoporins is a central driver of nuclear pore biogenesis. Cell Rep 44, 115468 (2025).

59. North, B.J., Fracchiolla, D., Ragusa, M.J., Martens, S. & Shoemaker, C.J. The rapidly expanding role of LC3-interacting regions in autophagy. J Cell Biol 224 (2025).

60. Ichimura, Y. et al. Structural basis for sorting mechanism of p62 in selective autophagy. J Biol Chem 283, 22847–22857 (2008).

61. Noda, N.N. et al. Structural basis of target recognition by Atg8/LC3 during selective autophagy. Genes Cells 13, 1211–1218 (2008).

62. Noda, N.N., Ohsumi, Y. & Inagaki, F. Atg8-family interacting motif crucial for selective autophagy. FEBS Lett 584, 1379–1385 (2010).

63. Johansen, T. & Lamark, T. Selective autophagy mediated by autophagic adapter proteins. Autophagy 7, 279–296 (2011).

64. Sahoo, M.R. et al. Nup358 binds to AGO proteins through its SUMO-interacting motifs and promotes the association of target mRNA with miRISC. EMBO Rep 18, 241–263 (2017).

65. Hamada, M. et al. Ran-dependent docking of importin-beta to RanBP2/Nup358 filaments is essential for protein import and cell viability. J Cell Biol 194, 597–612 (2011).

66. D’Acunzo, P. et al. Reversible induction of mitophagy by an optogenetic bimodular system. Nat Commun 10, 1533 (2019).

67. Xian, H., Yang, Q., Xiao, L., Shen, H.M. & Liou, Y.C. STX17 dynamically regulated by Fis1 induces mitophagy via hierarchical macroautophagic mechanism. Nat Commun 10, 2059 (2019).

68. McLelland, G.L., Lee, S.A., McBride, H.M. & Fon, E.A. Syntaxin-17 delivers PINK1/parkin-dependent mitochondrial vesicles to the endolysosomal system. J Cell Biol 214, 275–291 (2016).

69. Yang, A. et al. Autophagy Sustains Pancreatic Cancer Growth through Both Cell-Autonomous and Nonautonomous Mechanisms. Cancer Discov 8, 276–287 (2018).

70. Towers, C.G. et al. Cancer Cells Upregulate NRF2 Signaling to Adapt to Autophagy Inhibition. Dev Cell 50, 690–703 e696 (2019).

71. Ge, Y. et al. Enzyme-Mediated Intercellular Proximity Labeling for Detecting Cell-Cell Interactions. J Am Chem Soc 141, 1833–1837 (2019).

72. Niopek, D., Wehler, P., Roensch, J., Eils, R. & Di Ventura, B. Optogenetic control of nuclear protein export. Nat Commun 7, 10624 (2016).

73. Lam, A.J. et al. Improving FRET dynamic range with bright green and red fluorescent proteins. Nat Methods 9, 1005–1012 (2012).

74. Agrotis, A., Pengo, N., Burden, J.J. & Ketteler, R. Redundancy of human ATG4 protease isoforms in autophagy and LC3/GABARAP processing revealed in cells. Autophagy 15, 976–997 (2019).

