## Supplementary figures and images for "Rapid optogenetic blockade of autophagy reveals that nuclear pore complex proteins are robust autophagy substrates"

### Extended Data Fig. 1

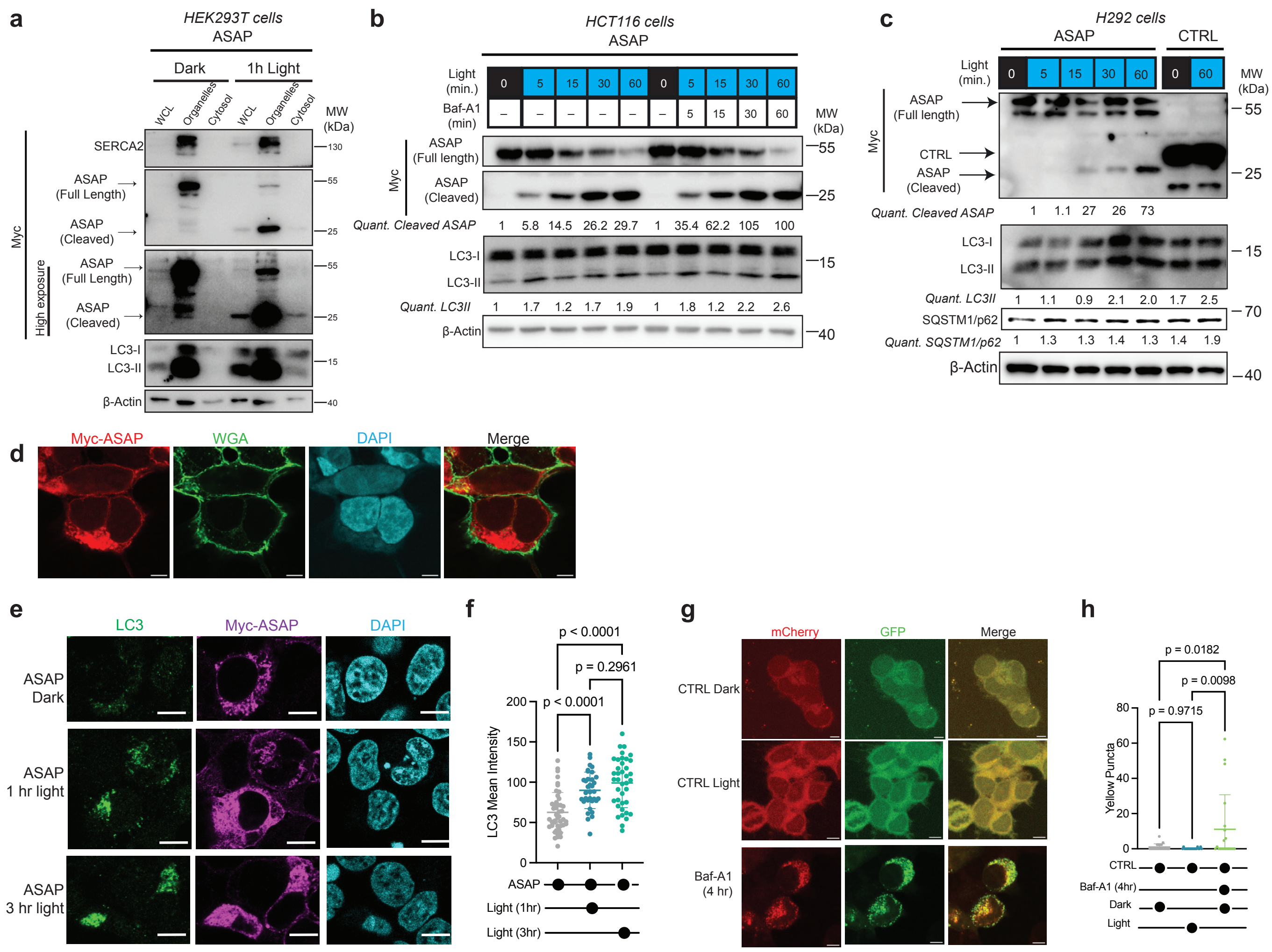

### Extended Data Fig. 2

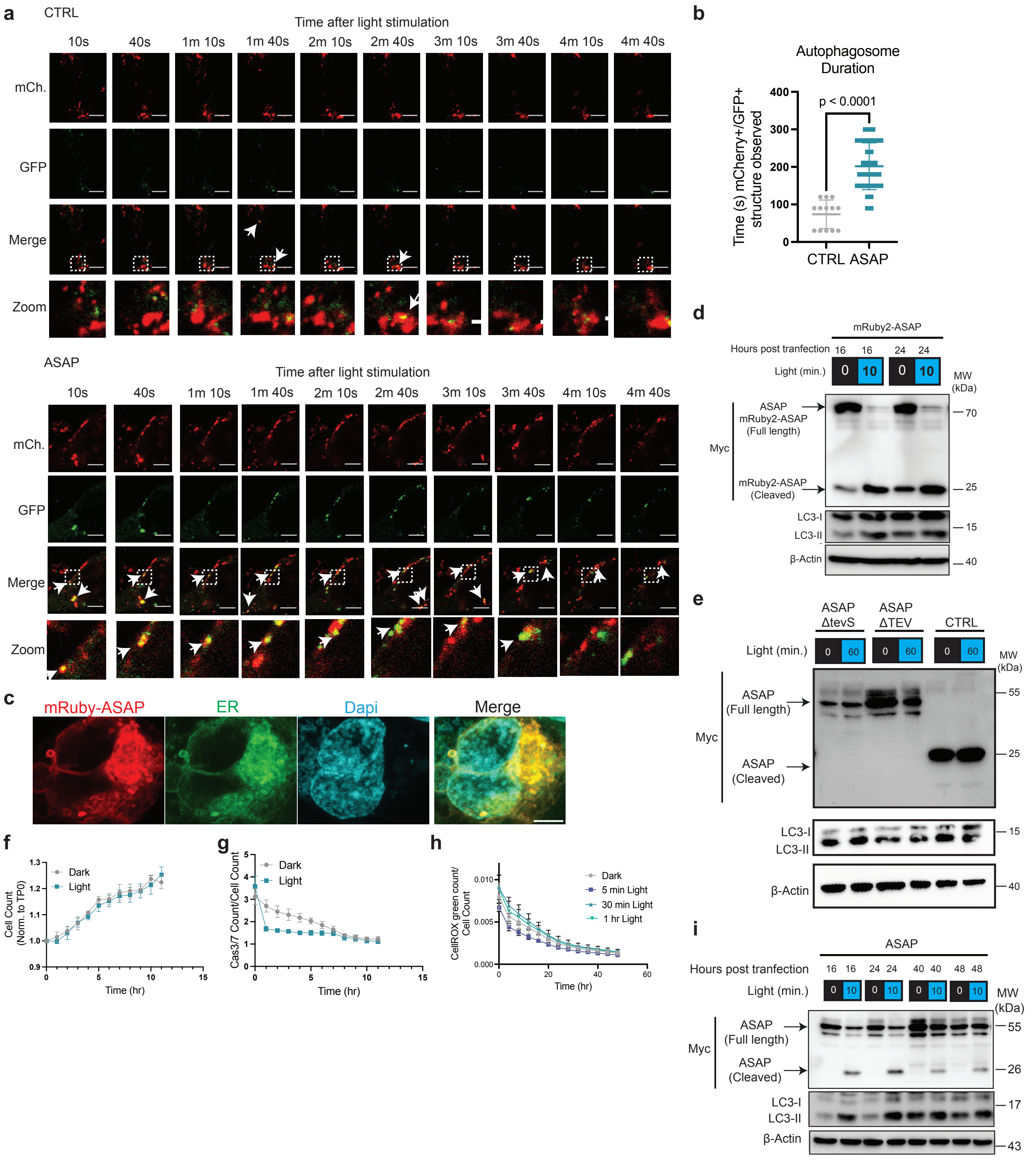

### Extended Data Fig. 3

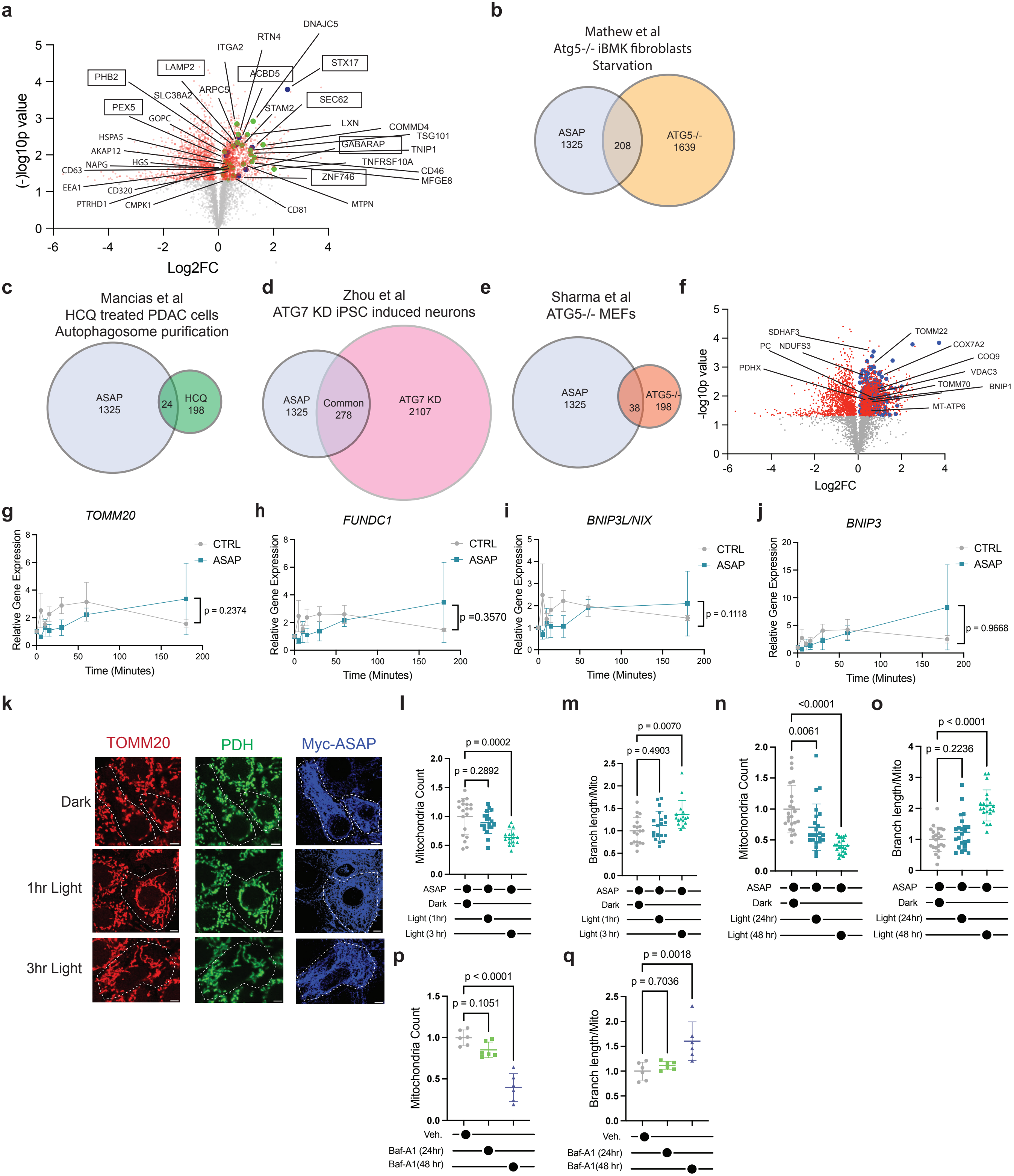

### Extended Data Fig. 4

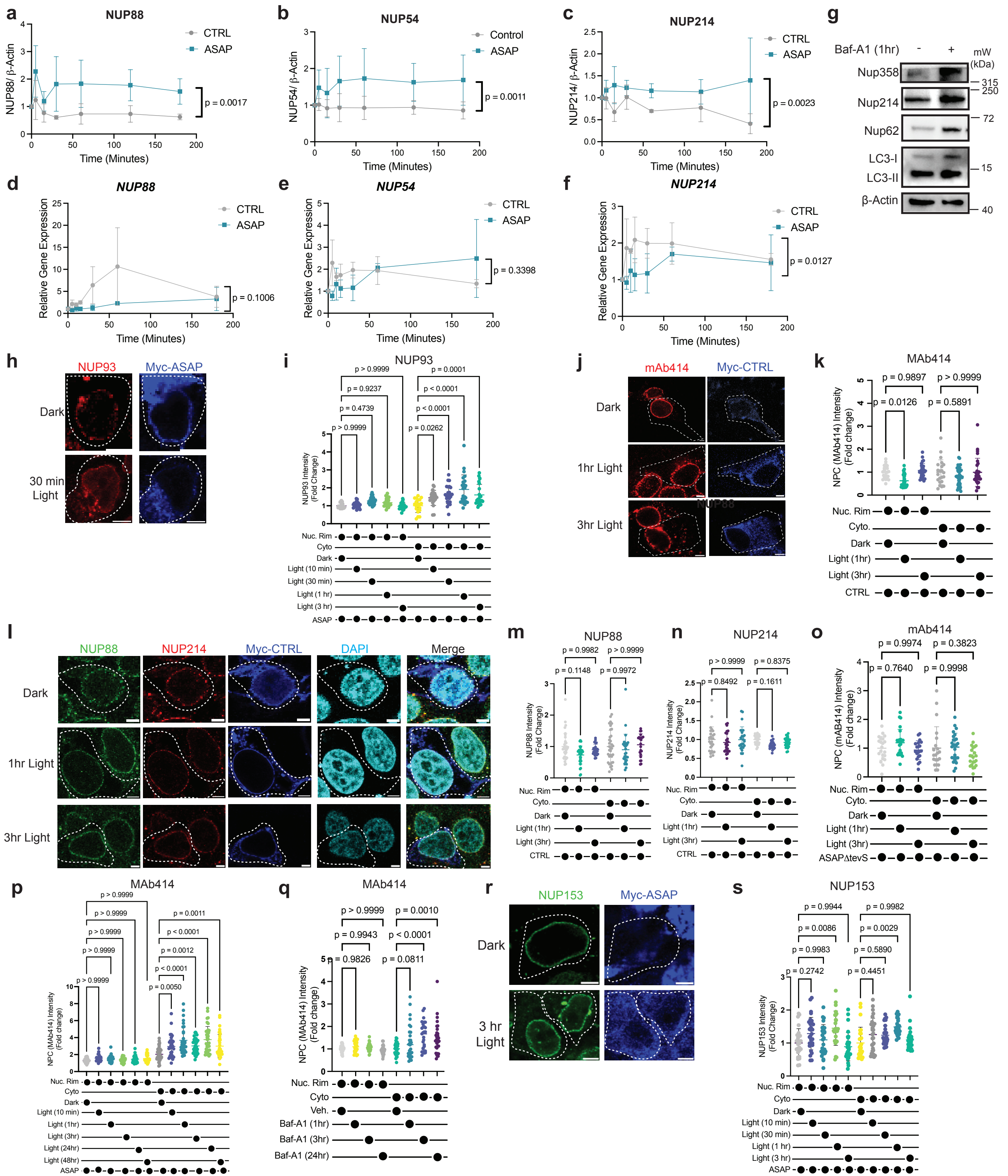

### Extended Data Fig. 5

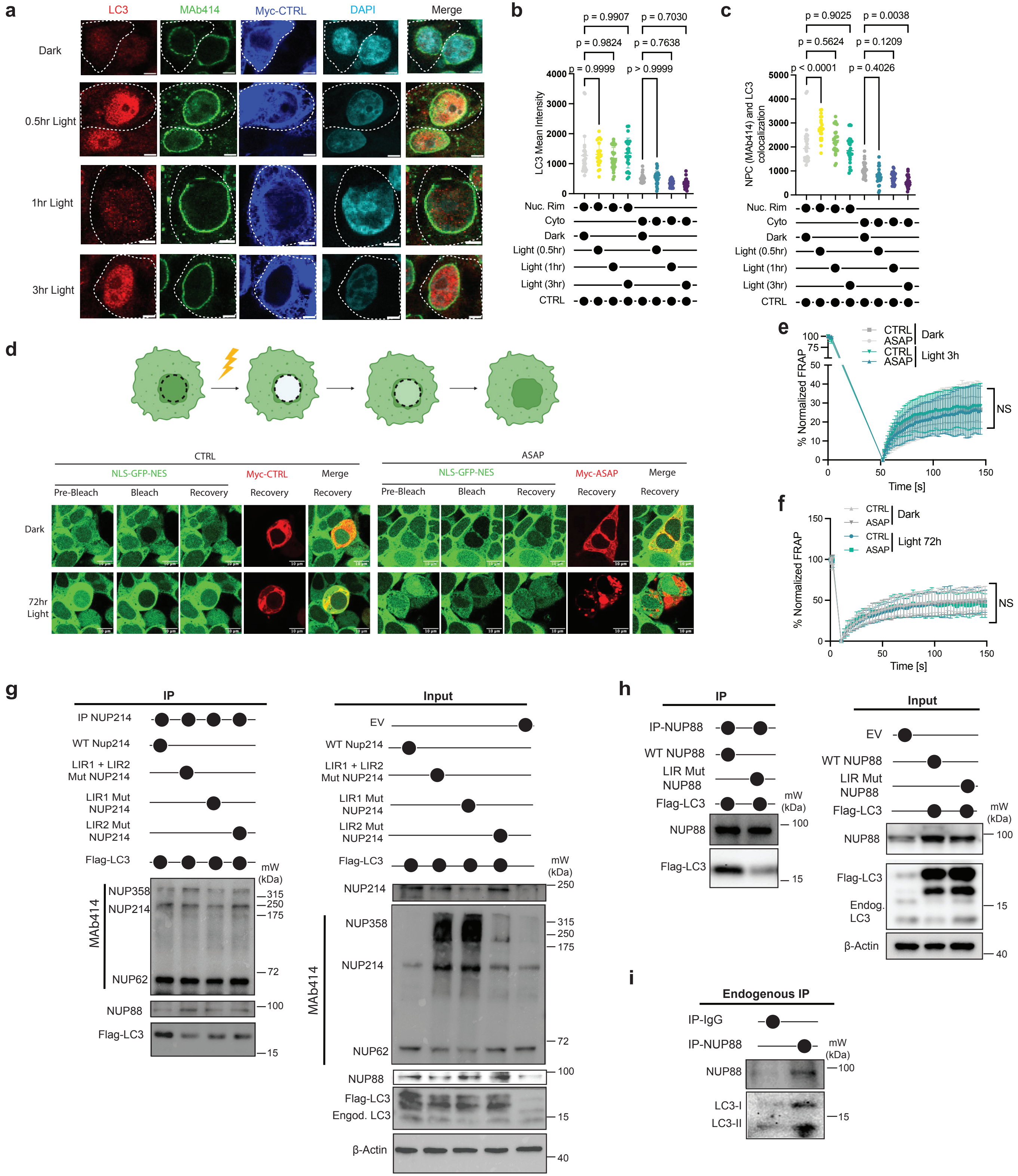

### Extended Data Fig. 6

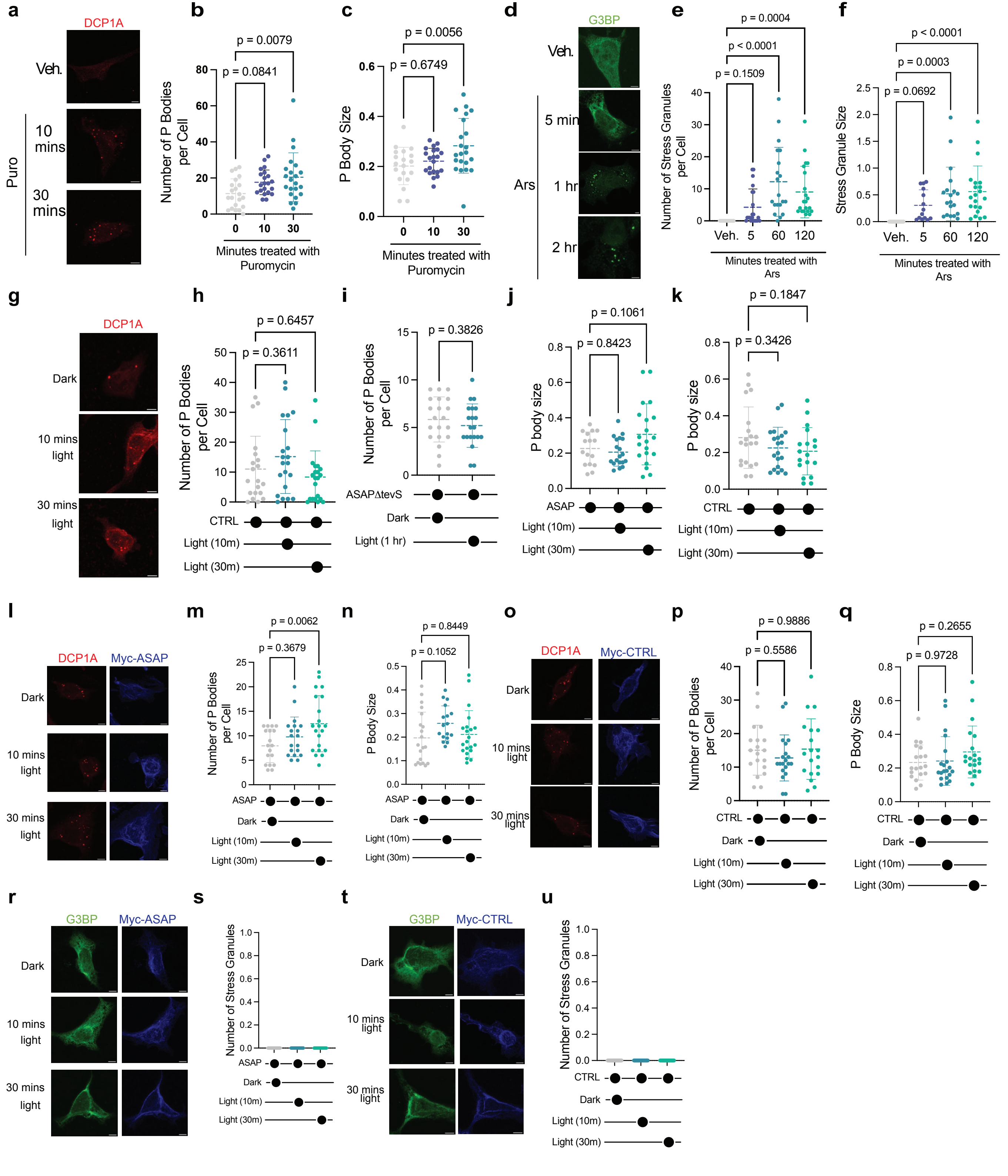

### Extended Data Fig. 7

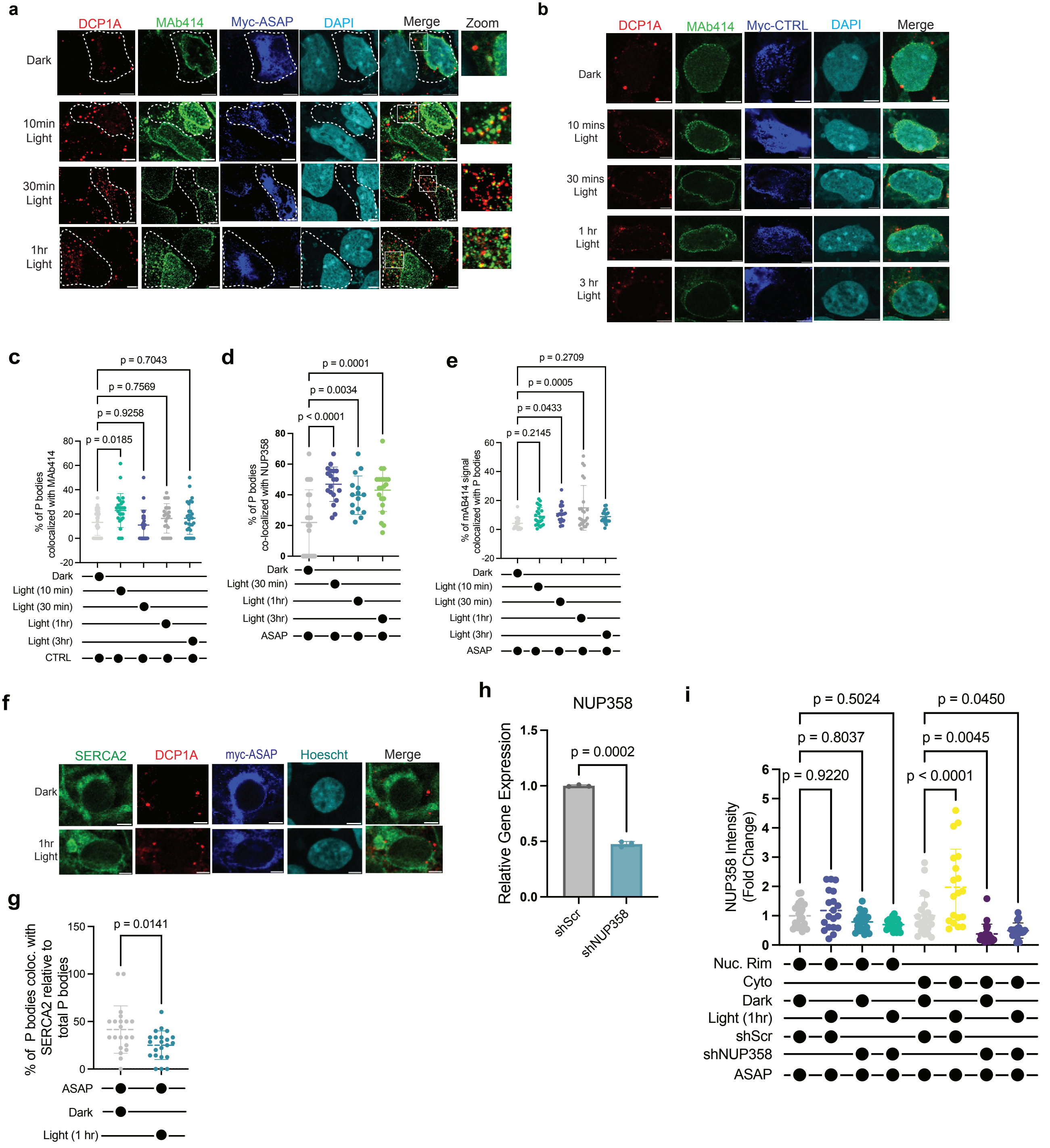
