## Extended Data Fig. 8 for "Rapid optogenetic blockade of autophagy reveals that nuclear pore complex proteins are robust autophagy substrates"

**a**

**ASAP**

METDTLLLWVLLLWVPGSTGDGGSNAVGQDTQEVIVVPHSLPFKVVVISAILALVVLTII
SLIILIMLWQKKPRCTDSAGSAGSAGELAEKLAGLDINGGASGSRATTLERIEKSFVIT
DPRLPDNPIIFVSDSFLQLTEYSREEILGRNCRFLQGPETDRATVRKIRDAIDNQTEVTV
QLINYTKSGKKFWNVFHLQPMRDYKGDVQYFIGVQLDGTERLHGAAEREAVCLVKK
TAFQIAENLYFQGGSGSGEQKLISEEDLNGEQKLISEEDLNGEQKLISEEDLNARELG
NSRVRETDPIPQDQNAAESWETLEADLIELSQLVTDFSLLVNSQQEKIDSIADHVNSA
AVNVEEGTKNLGKAAKYKLAALPVAGALIGGMVGGPIGLLAGFKVAGIAAALGGGVL
GFTGGKLIQRKKQKMMEKLTSSCPDLPSQTDKKCSTGGGSGSGSGSGATNFSLLKQ
AGDVEENPGPGSGSGSGSGSGSGSGESLFKGPRDYNPISSTICHLTNESDGHTTSLY
GIGFGPFIITNKHLFRRNNGTLLVQSLHGVFKVKNTTTLQQHLIDGRDMIIIRMPKDFPP
FPQKLKFREPQREERICLVTTNFQTKSMSSMVSDTSCTFPSSDGIFWKHWIQTKDGQ
CGSPLVSTRDGFIVGIHSASNFTNTNNYFTSVPKNFMELLTNQEAQQWVSGWRLNAD
SVLWGGHKVFMVYPYDVPDYA*

IgK leader sequence PDGFR TM hLOV1 tevS 3x Myc DN-STX17 P2A TEV HA

**b**

**ASAP-mRuby2**

METDTLLLWVLLLWVPGSTGDGGSNAVGQDTQEVIVVPHSLPFKVVVISAILALVVLTII
SLIILIMLWQKKPRCTDSAGSAGSAGVSKGEELIKENMRMKVVMEGSVNGHQFKCTG
EGEGNPYMGTQTMRIKVIEGGPLPFAFDILATSFMYGSRTFIKYPKGIPDFFKQSFPEG
FTWERVTRYEDGGVVTVMQDTSLEDGCLVYHVQVRGVNFPSNGPVMQKKTKGWEP
NTEMMYPADGGLRGYTHMALKVDGGGHLSCSFVTTYRSKKTVGNIKMPGIHAVDHR
LERLEESDNEMFVVQREHAVAKFAGLGGGMDELYKELAEKLAGLDINGGASGSRAT
TLERIEKSFVITDPRLPDNPIIFVSDSFLQLTEYSREEILGRNCRFLQGPETDRATVRKIR
DAIDNQTEVTVQLINYTKSGKKFWNVFHLQPMRDYKGDVQYFIGVQLDGTERLHGAA
EREAVCLVKKTAFQIAENLYFQGGSGSGEQKLISEEDLNGEQKLISEEDLNGEQKLISE
EDLNARELGNSRVRETDPIPQDQNAAESWETLEADLIELSQLVTDFSLLVNSQQEKID
SIADHVNSAAVNVEEGTKNLGKAAKYKLAALPVAGALIGGMVGGPIGLLAGFKVAGIA
AALGGGVLGFTGGKLIQRKKQKMMEKLTSSCPDLPSQTDKKCSTGGGSGSGSGSG
ATNFSLLKQAGDVEENPGPGSGSGSGSGSGSGSGESLFKGPRDYNPISSTICHLTNE
SDGHTTSLYGIGFGPFIITNKHLFRRNNGTLLVQSLHGVFKVKNTTTLQQHLIDGRDMIII
RMPKDFPPFPQKLKFREPQREERICLVTTNFQTKSMSSMVSDTSCTFPSSDGIFWKH
WIQTKDGQCGSPLVSTRDGFIVGIHSASNFTNTNNYFTSVPKNFMELLTNQEAQQWV
SGWRLNADSVLWGGHKVFMVYPYDVPDYA*

IgK leader sequence PDGFR TM mRuby2 hLOV1 tevS 3x Myc DN-STX17 P2A TEV HA

**c**

**Control**

METDTLLLWVLLLWVPGSTGDGGSNAVGQDTQEVIVVPHSLPFKVVVISAILALVVLTII
SLIILIMLWQKKPRCTDSAGSAGSAGSRATTLERIEKSFVITDPRLPDNPIIFVSDSFLQL
TEYSREEILGRNCRFLQGPETDRATVRKIRDAIDNQTEVTVQLINYTKSGKKFWNLFH
LQPMRDQKGDVQYFIGVQLDGTERVRDAAEREAVMLVKKTAEEIDEAAKENLYFQM
GSGSGEQKLISEEDLNGEQKLISEEDLNGEQKLISEEDL

IgK leader sequence PDGFR TM eLOV tevS 3x Myc

**d**

**ASAPΔTEV**

METDTLLLWVLLLWVPGSTGDGGSNAVGQDTQEVIVVPHSLPFKVVVISAILALVVLTII
SLIILIMLWQKKPRCTDSAGSAGSAGELAEKLAGLDINGGASGSRATTLERIEKSFVIT
DPRLPDNPIIFVSDSFLQLTEYSREEILGRNCRFLQGPETDRATVRKIRDAIDNQTEVTV
QLINYTKSGKKFWNVFHLQPMRDYKGDVQYFIGVQLDGTERLHGAAEREAVCLVKK
TAFQIAENLYFQGGSGSGEQKLISEEDLNGEQKLISEEDLNGEQKLISEEDLNARELG
NSRVRETDPIPQDQNAAESWETLEADLIELSQLVTDFSLLVNSQQEKIDSIADHVNSA
AVNVEEGTKNLGKAAKYKLAALPVAGALIGGMVGGPIGLLAGFKVAGIAAALGGGVL
GFTGGKLIQRKKQKMMEKLTSSCPDLPSQTDKKCSTGGGSGSGSGSGATNFSLLKQ
AGDVEENPGPGSGSGSGSGSGSGSYPYDVPDYA*

IgK leader sequence PDGFR TM hLOV1 tevS 3x Myc DN-STX17 P2A HA

**e**

**ASAPΔtevS**

METDTLLLWVLLLWVPGSTGDGGSNAVGQDTQEVIVVPHSLPFKVVVISAILALVVLTII
SLIILIMLWQKKPRCTDSAGSAGSAGELAEKLAGLDINGGASGSRATTLERIEKSFVIT
DPRLPDNPIIFVSDSFLQLTEYSREEILGRNCRFLQGPETDRATVRKIRDAIDNQTEVTV
QLINYTKSGKKFWNVFHLQPMRDYKGDVQYFIGVQLDGTERLHGAAEREAVCLVKK
TAFQIAGSGSGEQKLISEEDLNGEQKLISEEDLNGEQKLISEEDLNARELGNSRVRET
DPIPQDQNAAESWETLEADLIELSQLVTDFSLLVNSQQEKIDSIADHVNSAAVNVEEGT
KNLGKAAKYKLAALPVAGALIGGMVGGPIGLLAGFKVAGIAAALGGGVLGFTGGKLI
QRKKQKMMEKLTSSCPDLPSQTDKKCSTGGGSGSGSGSGATNFSLLKQAGDVEEN
PGPGSGSGSGSGSGSGSGESLFKGPRDYNPISSTICHLTNESDGHTTSLYGIGFGPFII
TNKHLFRRNNGTLLVQSLHGVFKVKNTTTLQQHLIDGRDMIIIRMPKDFPPFPQKLKF
REPQREERICLVTTNFQTKSMSSMVSDTSCTFPSSDGIFWKHWIQTKDGQCGSPLVS
TRDGFIVGIHSASNFTNTNNYFTSVPKNFMELLTNQEAQQWVSGWRLNADSVLWGG
HKVFMVYPYDVPDYA*

IgK leader sequence PDGFR TM hLOV1 3x Myc DN-STX17 P2A TEV HA

**f**

**Control-mRuby2**

METDTLLLWVLLLWVPGSTGDGGSNAVGQDTQEVIVVPHSLPFKVVVISAILALVVLTII
SLIILIMLWQKKPRCTDSAGSAGSAGVSKGEELIKENMRMKVVMEGSVNGHQFKCTG
EGEGNPYMGTQTMRIKVIEGGPLPFAFDILATSFMYGSRTFIKYPKGIPDFFKQSFPEG
FTWERVTRYEDGGVVTVMQDTSLEDGCLVYHVQVRGVNFPSNGPVMQKKTKGWEP
NTEMMYPADGGLRGYTHMALKVDGGGHLSCSFVTTYRSKKTVGNIKMPGIHAVDHR
LERLEESDNEMFVVQREHAVAKFAGLGGGMDELYKSRATTLERIEKSFVITDPRLPDN
PIIFVSDSFLQLTEYSREEILGRNCRFLQGPETDRATVRKIRDAIDNQTEVTVQLINYTK
SGKKFWNLFHLQPMRDQKGDVQYFIGVQLDGTERVRDAAEREAVMLVKKTAEEIDE
AAKENLYFQMGSGSGEQKLISEEDLNGEQKLISEEDLNGEQKLISEEDL

IgK leader sequence PDGFR TM mRuby2 eLOV tevS 3x Myc

**g**

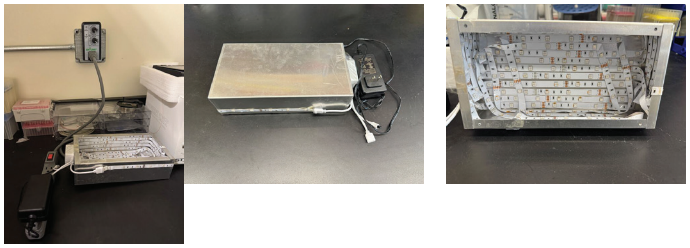

**h**

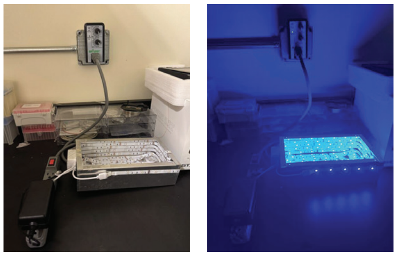

**i**

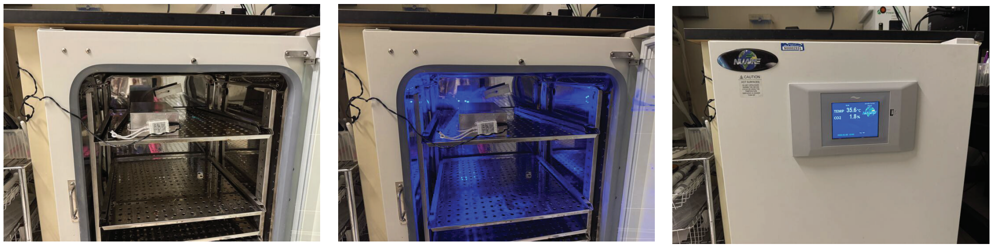

**j**

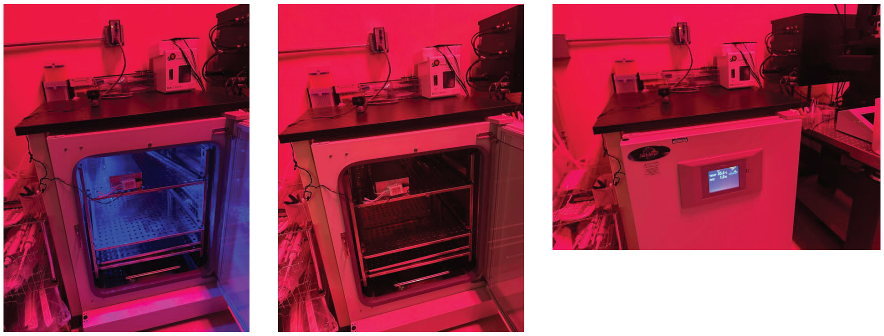
