## Extended Data Fig. 9 for "Rapid optogenetic blockade of autophagy reveals that nuclear pore complex proteins are robust autophagy substrates"

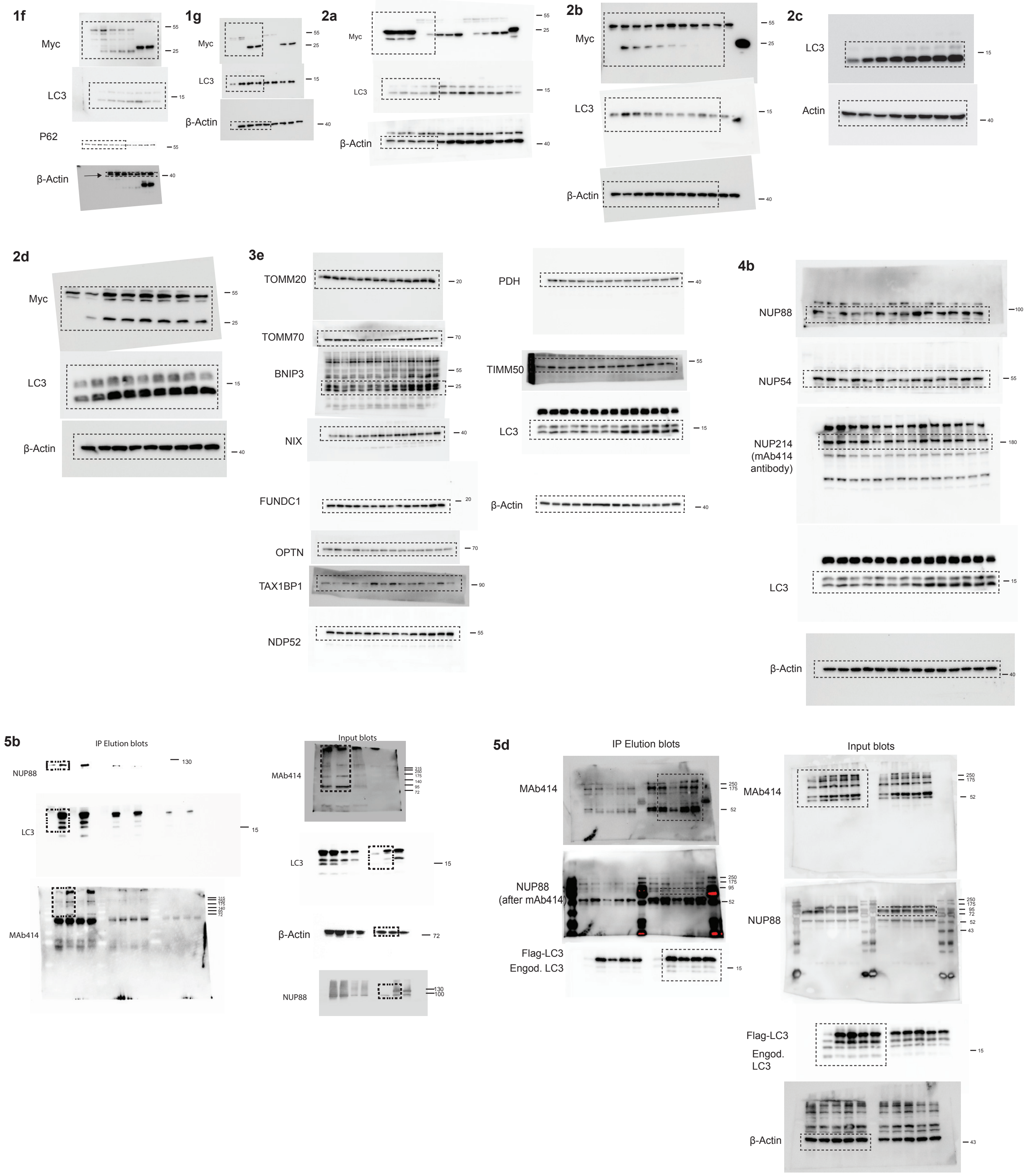

1f
Myc
LC3
P62
β-Actin
55
25
15
55
40
1g
Myc
LC3
β-Actin
55
25
15
40
2a
Myc
LC3
β-Actin
55
25
15
40
2b
Myc
LC3
β-Actin
55
25
15
40
2c
LC3
Actin
15
40
2d
Myc
LC3
β-Actin
55
25
15
40
3e
TOMM20
TOMM70
BNIP3
NIX
FUNDC1
OPTN
TAX1BP1
NDP52
20
70
55
25
40
20
70
90
55
PDH
TIMM50
LC3
β-Actin
40
55
15
40
4b
NUP88
NUP54
NUP214
(mAb414
antibody)
LC3
β-Actin
100
55
180
15
40
5b
IP Elution blots
NUP88
LC3
MAb414
130
15
315
250
175
140
95
72
Input blots
MAb414
LC3
β-Actin
NUP88
315
250
175
140
95
72
15
72
130
100
5d
IP Elution blots
MAb414
NUP88
(after mAb414)
Flag-LC3
Engod. LC3
250
175
52
250
175
95
52
15
Input blots
MAb414
NUP88
Flag-LC3
Engod.
LC3
β-Actin
250
175
52
250
175
95
72
52
43
15
43

**5e**

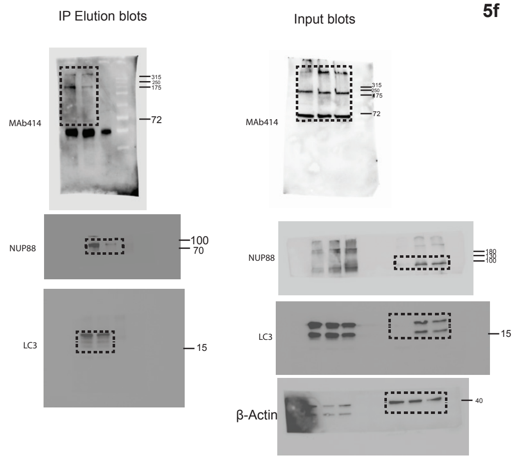


**5f**

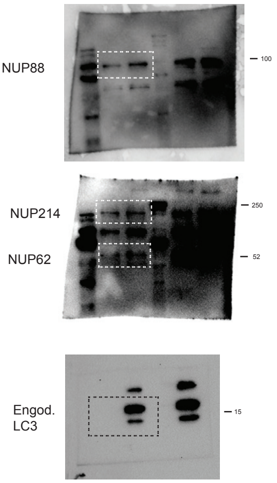


**Extended Data Fig. 1a**

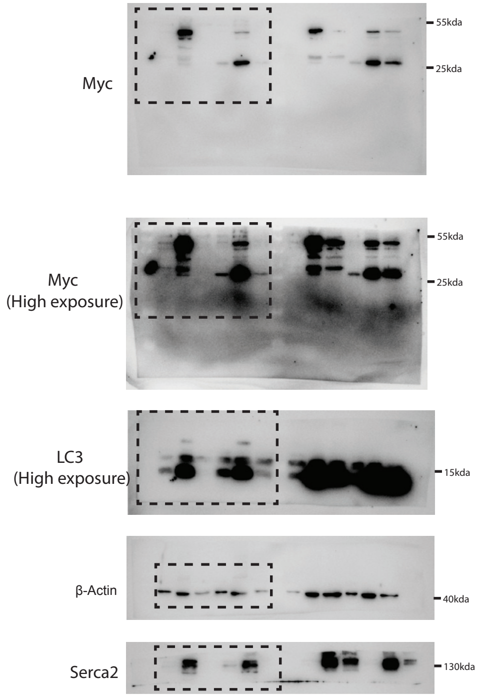


**Extended Data Fig. 1b**

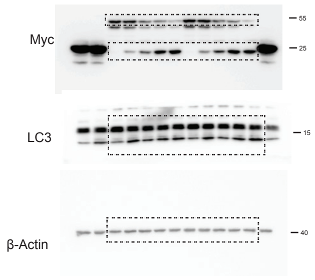


Extended Data Fig. 1c

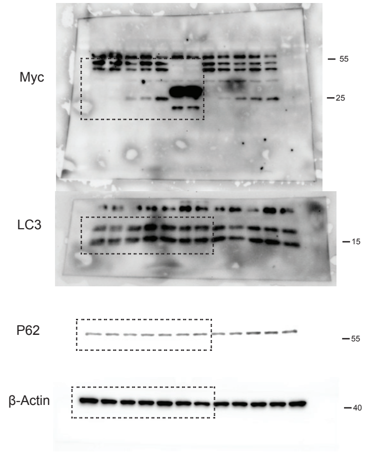


**Extended Data Fig. 2d**

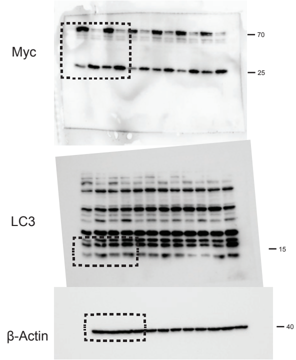


**Extended Data Fig. 2e**

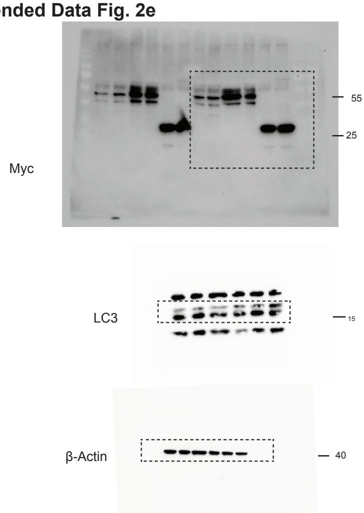


**Extended Data Fig. 2f**

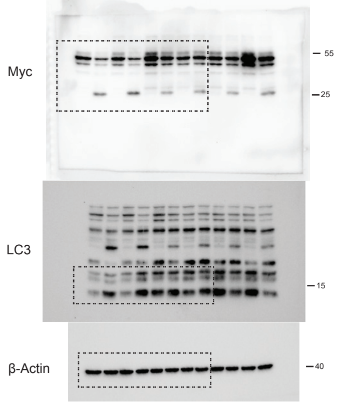

### Extended Data Fig. 4g

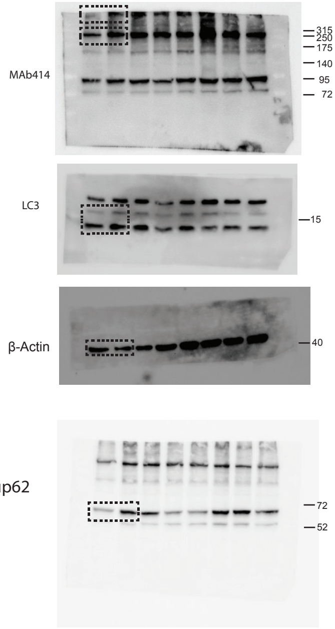


### Extended Data Fig. 5g

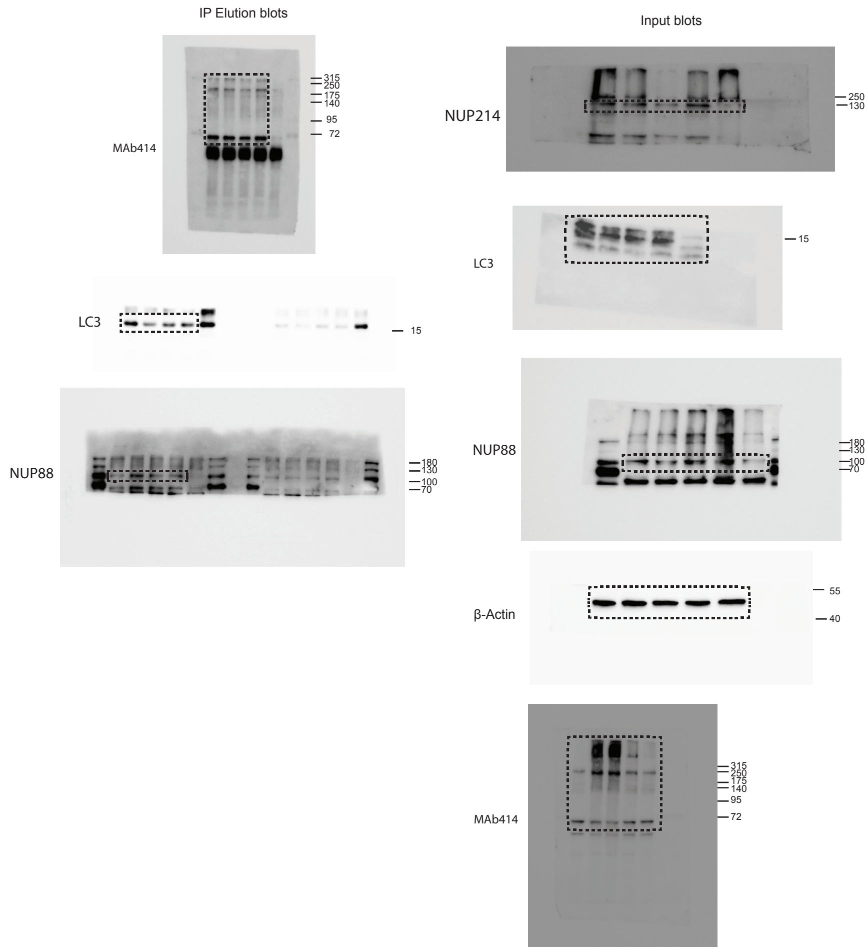


### Extended Data Fig. 5h

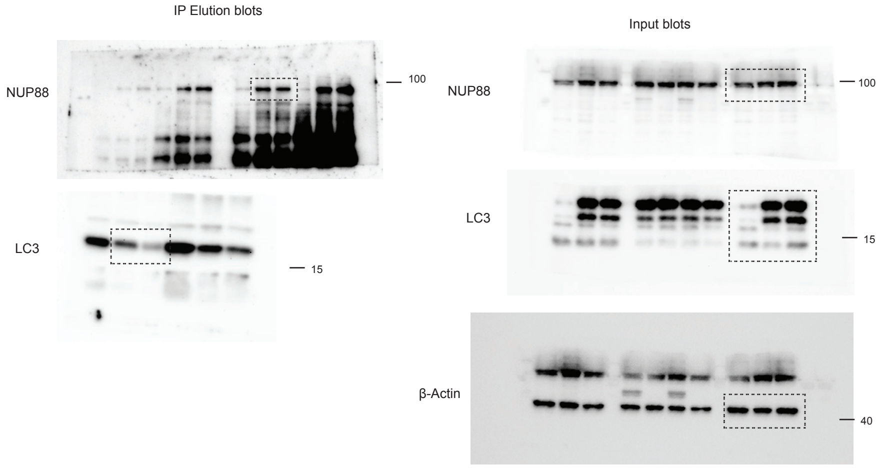


### Extended Data Fig. 5i

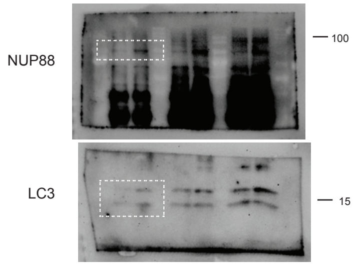
