## Extended Data Figure Legends for "Rapid optogenetic blockade of autophagy reveals that nuclear pore complex proteins are robust autophagy substrates"

**Extended Data** **Figure 1:** **ASAP inhibits autophagy within minutes in different cell lines. Related to Figure 1.**

(a) HEK293 cells transfected with ASAP and treated with 1hr of 450nm pulsed light. Western blot from whole cell lysates (WCL), organelle-rich fractions (Organelles), or organelle-depleted fractions (Cytosol) blotted for SERCA2 (ER marker), Myc indicating full length ASAP, cleaved ASAP, LC3, and β-Actin. Because autophagosomes are membrane bound vesicles, the fractionation cannot separate ER from autophagosomes. The organelle-rich fraction contains uncleaved ASAP in the dark which is significantly depleted in this fraction with 1hr of light (lane 2 vs. lane 5). This fraction also includes the cleaved ASAP with 1hr of light because autophagosomes are present in this fraction, indicated by the high levels of LC3-II. The cytosolic fraction is depleted of organelles including both ER and autophagosomes, and yet it still contains cleaved ASAP in the presence of light clearly showing a light induced dissociation from the ER. Blot is representative of 2 independent experiments.
(b) HCT116 cells transfected with ASAP and treated with indicated times of 450nm pulsed light with or without bafilomycin (Baf-A1; 10nM). Western blot for Myc indicating full length ASAP, cleaved ASAP, LC3, and β-Actin. Quantification of fold change in cleaved ASAP and LC3-II bands normalized β-Actin are indicated. Blot is representative of 3 independent experiments.

**(c)** NCI-H292 cells transfected with ASAP or CTRL and treated with indicated times of 450nm pulsed light. Western blot for Myc indicating full length ASAP, cleaved ASAP, and CTRL products, as well as SQSTM1/p62, LC3, and β-Actin. Quantification of fold change in cleaved ASAP, SQSTM1/p62, and LC3-II bands normalized β-Actin are indicated. Blot is representative of 2 independent experiments.

**(d**) Representative confocal images of HEK293T cells transfected with myc-ASAP and stained for Myc and WGA to mark the plasma membrane. Scale bar = 10μM. Data are representative of N = 2 experiments.

**(e-f)** HEK293T cells transfected with ASAP and treated with 450nm pulsed light for indicated periods of time followed by IF for LC3 and Myc-tagged ASAP**. (e)** Representative images, scale bar = 10μM. **(f)** Quantification of LC3 mean intensity in Myc-ASAP + cells. N = 34 – 45 cells per condition. Data are represented as mean ± S.D and are representative of N = 3 experiments. Statistical analyses were performed using an Ordinary one-way ANOVA.

**(g-h)** HCT116 cells with stable expression of mcherry-GFP-LC3 transfected with CTRL and treated with 450nm pulsed light for 5 minutes or with Baf-A1 for 4hrs (10nM) followed by **(g)** IF for mCherry and GFP puncta. **(h)** Quantification of yellow LC3 puncta. N = 38-65 cells per condition. Data are represented as mean ± S.D for 3-4 independent experiments. The CTRL Dark and CTRL light conditions displayed here are also displayed in Fig. 1J and reshown here to compare to BafA1 4hr treatment. Statistical analyses were performed using an Ordinary one-way ANOVA.

Uncropped western blots are provided in Extended Data Fig. 9

**Extended Data Figure** **2: ASAP, but not control constructs, block autophagy without light induced toxicity. Related to Figure 1 and 2.**

**(a-b)** HCT-116 cells with stable expression of mCherry-GFP-LC3 transfected with CTRL or ASAP stimulated with 450nm in a light box followed by live cell confocal imaging. Live imaging was started 10s later and Z stack images captured every 30s for approximately 5 minutes. Scale bars represent 5μM. (b) Quantification of autophagosome duration, measured as the amount of time (seconds) that a mCherry+/GFP+ structure is observed. Statistical analyses were performed using a two-tailed T-test.

**(c)** Live cell confocal imaging in HEK293T cells transfected with mRuby2-ASAP (red) and the ER marker, ER-Tracker Green. Scale bars represent 5μM. Images are representative of 2 independent experiments.

**(d)** HEK293T cells transfected with mRuby2-ASAP and left in the dark for indicated periods of time followed by a 10 minute light treatment with 450nm pulsed light. Western blot for Myc indicating full length mRuby-ASAP, cleaved ASAP, as well as LC3, and β-Actin. Blot is representative of N = 2 experiments.

(**e**) HEK293T cells transfected with ASAPΔtevS, ASAPΔTEV, or CTRL and treated with 1hr of 450nm pulsed light. Western blot for Myc indicating full length ASAP, cleaved ASAP, and CTRL products, as well as LC3, and β-Actin. Blot is representative of N = 2 experiments.

**(f-g)** Incucyte live cell imaging in HCT116 cells treated with pulsed light every 30 minutes for indicated time courses. **(f)** Fold change in cell count based on nuc-red staining. Data graphed as mean ± S.D for technical replicates of 5 wells per condition and is representative of 2 independent experiments. **(g)** Quantification of Cell Event Caspase 3/7 normalized to cell count (based on nuc-red). Data graphed as mean ± S.D for technical replicates of 5 wells per condition and is representative of 2 independent experiments.

**(h)** Incucyte live cell imaging in HCT116 cells treated with pulsed light for indicated periods of time and then placed back in the dark for the remainder of the time course. Quantification of Cell ROX green count normalized to cell count (based on nuc-red). Data graphed as mean ± S.D for technical replicates of 5 wells per condition and is representative of 2 independent experiments.

(**i**) HEK293T cells transfected with ASAP and left in the dark for indicated periods of time followed by 10 min treatment of 450nm pulsed light. Western blot for Myc indicating full length ASAP and cleaved ASAP, well as LC3, and β-Actin. Blot is representative of N = 2 experiments.

Uncropped western blots are provided in Extended Data Fig. 9 **Extended Data Figure** **3:** **TMT proteomics with ASAP confirms known autophagy substrates and identifies unique autophagy substrates. Related to Figure 3.**

**(a)** Volcano plot depicting Log_2_ fold change and (-)Log_10_ P value for proteins altered in ASAP expressing cells versus CTRL cells after 30 minutes of 450nm pulsed light. Proteins with a p value < 0.05 are indicated in red. ATG8 family proteins, STX17 and known autophagy substrates are highlighted in green and blue where the blue proteins are the same proteins highlighted in Figure 3c. Additional proteins are highlighted here that were previously identified from other proteomics studies that assess autophagy substrates in different systems and stress-induced conditions^16, 19-21^.

**(b-e)** Venn diagram showing overlapping and unique proteins that accumulate with autophagy inhibition from other proteomics studies including **(b)** Mathew et al^16^, **(c)** Mancias et al^19^, **(d)** Zhou et al^21^, and **(e)** Sharma et al^20^.

**(f)** Volcano plot depicting Log_2_ fold change and (-)Log_10_ P value for proteins altered in ASAP expressing cells versus CTRL cells after 30 minutes of 450nm pulsed light. Proteins with a p value < 0.05 are indicated in red. Data is the same as what is represented in Figure 2c. Mitochondrial proteins that accumulate with light are highlighted in blue.

**(g-j)** HEK293T cells transfected with CTRL or ASAP and treated with 450nm pulsed light for indicated periods of time followed by qRT-PCR for gene expression of indicated genes. Fold change in gene expression relative to the respective Dark conditions and normalized to housekeeping gene, *18s,* for **(g)** *TOMM20*, **(h)** *FUNDC1*, **(i)** *BNIP3L/NIX*, and **(j)** *BNIP3*. Data shown as the average ± SEM for 3 independent experiments. Statistical analysis was performed using two-tailed T-test between ASAP and CTRL based on all time points.

**(k-o)** NCI-H292 cells transfected with ASAP and treated with 450nm pulsed light for indicated periods of time followed by IF for TOMM20 and Myc. **(k)** Representative IF images of TOMM20, PDH, and Myc-ASAP. Dotted white lines indicate examples of ASAP positive cells. Scale bar = 5μM. Quantification of **(l, n)** the fold change in mitochondrial count based on TOMM20 signal and **(m, o)** the fold change in mitochondrial branch length using Mitochondria Analyzer Plugin on ImageJ. N = 15-23 cells per condition. Data are graphed as mean ± S.D. and are representative of 2 independent experiments. Statistical analyses were performed using a One Way ANOVA.

(**p-q**) NCI-H292 cells treated with Veh. (DMSO) or Baf-A(10nM) for indicated periods of time followed by IF for TOMM20. Quantification of **(p)** the fold change in mitochondrial count based on TOMM20 signal and **(q)** the fold change in mitochondrial branch length using Mitochondria Analyzer Plugin on ImageJ. N = 6 images per condition, each containing 5-10 cells. Data are graphed as mean ± S.D. and are representative of 2 independent experiments. Statistical analyses were performed using a One Way ANOVA

**Extended Data Figure** **4:** **Cytoplasmic nuclear pore complex proteins accumulate after rapid autophagy inhibition in ASAP but not CTRL expressing cells. Related to Figure 4.**

**(a-c)** HEK293T cells transfected with ASAP or CTRL and treated with 450nm pulsed light for indicated periods of time. Quantification of western blot bands shown in Fig. 4b for **(a)** NUP88, **(b)** NUP54, and **(c)** NUP214 are normalized to β-Actin and graphed as the fold change to the respective dark conditions for ASAP and CTRL expressing cells. Data shown as the average ± SEM for N = 3 experiments. The area under the curve from these graphs is shown in Figure 4c. Statistical analysis was performed using two-tailed T-test between ASAP and CTRL taking into account all time points.

**(d-f)** HEK293T cells transfected with CTRL or ASAP and treated with 450nm pulsed light for indicated periods of time followed by qRT-PCR for gene expression of indicated genes. Fold change in gene expression relative to the respective Dark conditions and normalized to housekeeping gene, *18s,* for **(d)** *NUP88*, **(e)** *NUP54*, and **(f)** *NUP214*. Data shown as the average ± SEM for 3 independent experiments. Statistical analysis was performed using two-tailed T-test between ASAP and CTRL.

**(g)** HEK293T cells treated with Baf-A1 for 1hr (10nM) followed by immunoblotting for MAb414 and bands are shown for NUP358, NUP214, NUP62, and LC3. Western blot is representative of 3 independent experiments.

**(h-n)** HEK293T cells transfected with CTRL or ASAP as indicated and treated with 450nm pulsed light for indicated periods of time followed by IF. **(h,j,l)** Representative IF images of the pan NPC antibody (MAb414), NUP88, NUP214, NUP93 and Myc-tag for ASAP. Dotted white lines indicate examples of ASAP positive cells. Scale bar = 5μM. **(i,k.m,n)** Quantification in the fold change of nucleoporin signal that is cytoplasmic or on the nuclear rim normalized to the respective Dark condition. N=21-37 cells per condition. Data are graphed as mean ± S.D. and are representative of 2-3 independent experiments. Statistical analyses were performed using a One Way ANOVA.

(**o**) HEK293T cells transfected with ASAPΔtevS and treated with 450nm pulsed light for indicated periods of time followed by IF for MAb414 and Myc. Quantification in the fold change of nucleoporin signal that is cytoplasmic or on the nuclear rim normalized to the respective Dark condition. N=21-25 cells per condition. Data are graphed as mean ± S.D. and are representative of 2 independent experiments. Statistical analyses were performed using a One Way ANOVA.

(**p**) HEK293T cells transfected with ASAP and treated with 450nm pulsed light for indicated periods of time followed by IF for MAb414 and Myc. For time points that are 1hr or less, pulsed light is continuous. For time points greater than 1hr, the cells are treated with 5 minutes of pulsed light every 30 minutes. Quantification in the fold change of nucleoporin signal that is cytoplasmic or on the nuclear rim normalized to the respective Dark condition. N=24-46 cells per condition. Data are graphed as mean ± S.D. and are representative of 2 independent experiments. Statistical analyses were performed using a One Way ANOVA.

**(q**) HEK293T cells treated with Baf-A1 (10nM) for indicated periods of time followed by IF for MAb414. Quantification in the fold change of nucleoporin signal that is cytoplasmic or on the nuclear rim normalized to Veh. treated conditions. N=28-32 cells per condition. Data are graphed as mean ± S.D. and are representative of 2-3 independent experiments. Statistical analyses were performed using a One Way ANOVA.

(**q-r**) HEK293T cells transfected with ASAP and treated with 450nm pulsed light for indicated periods of time followed by IF. **(q)** Representative IF images of NUP153 and Myc-tag for ASAP. Dotted white lines indicate examples of ASAP positive cells. Scale bar = 5μM. **(r)** Quantification in the fold change of nucleoporin signal that is cytoplasmic or on the nuclear rim normalized to the respective Dark condition. N=24-29 cells per condition. Data are graphed as mean ± S.D. and are representative of 2 independent experiments. Statistical analyses were performed using a One Way ANOVA.

Uncropped western blots are provided in Extended Data Fig. 9

**Extended Data Figure** **5: NPC function is not altered with ASAP and NUP214 and NUP88 have bona fide LC3 interacting regions. Related to Figures 4 and 5.**

**(a-c)** HEK293T cells transfected with CTRL and treated with 450nm pulsed light for indicated periods of time followed by IF. **(a)** Representative IF images of the MAb414, Myc-CTRL, LC3, and Dapi. Dotted white lines indicate examples of ASAP positive cells. Scale bar = 5μM. Quantification of **(b)** LC3 intensity and **(c)** MAb414 colocalization with LC3 on the nuclear rim or in the cytoplasm. N = 24-31 cells per condition. Data are graphed as mean ± S.D. and are representative of 3 independent experiments. Statistical analyses were performed using a One Way ANOVA.

**(d-f)** HEK293T cells stably expressing the transport reporter NLS-2xGFP-NES and transfected with ASAP-mRuby2 or CTRL-mRuby2 and treated with 450nm pulsed light (**d**) Top: Animation of how FRAP assay works to measure NPC function by assessing the nuclear signal before and after bleaching to measure signal recovery indicative of transport through the NPC. Bottom: Representative images of NLS-GFP-NES pre and post bleach in cells treated with pulsed light for 72hrs. Quantification of nuclear GFP signal in cells treated with (**e**) 3hrs of light or (**f**) 72hrs of light. Images obtained every 3 seconds. N = 15-21 cells per condition. Data are graphed as mean ± S.D. and are representative of 2 independent experiments. Statistical analyses were performed using a One Way ANOVA.

**(g)** HEK293T cells expressing Flag-LC3 and either WT GFP-NUP214 or GFP-NUP214 with LIR mutants. IP of NUP214 followed by immunoblotting for NUP214, MAb414, and LC3. (Left) Blots from IP elution and (right) blots from the input samples prior to IP. Blots are representative of 2 independent experiments.

**(h)** HEK293T cells expressing Flag-LC3 and either WT or NUP88 mutants. IP of NUP88 followed by immunoblotting for NUP88 and LC3. (Left) Blots from IP elution and (right) blots from the input samples prior to IP. Blots are representative of 2 independent experiments

**(i)** Endogenous IP of NUP88 in HEK293T cells, followed by immunoblotting for LC3 and NUP88. Blots are representative of 2 independent experiments.

Uncropped western blots are provided in Extended Data Fig. 9

**Extended Data Figure 6:** **Rapid autophagy inhibition induces P bodies but has no effect on stress granules. Related to Figure 6.**

**(a-c)** HEK293T cells treated with Puromycin (100 μg/mL) for indicated periods of time followed by IF for DCP1A to mark P bodies. **(a)** Representative images. **(b)** Quantification of the number of P bodies per cell. **(c)** Quantification of the size of P bodies in pixels. N = 20-21 cells per condition. Data are graphed as mean ± S.D. and are representative of 2 independent experiments. Statistical analyses were performed using a One Way ANOVA.

**(d-f)** HEK293T cells treated with Sodium Arsenite (Ars, 100 μg/mL) for indicated periods of time followed by IF for G3BP to mark stress granules. **(d)** Representative images. **(e)** Quantification of the number of stress granules per cell. **(f)** Quantification of the size of stress granules in pixels. N = 14-21 cells per condition. Data are graphed as mean ± S.D. and are representative of 2 independent experiments. Statistical analyses were performed using a One Way ANOVA.

**(g-i)** HEK293T cells transfected with CTRL or ASAPΔtevs and treated with 450nm pulsed light for indicated periods of time followed by IF. **(g)** Representative IF images of the P body marker, DCP1A. Scale bar = 5μM. **(h)** Quantification of the number of P bodies per cell in cells expressing CTRL with indicated periods of light. N=20 cells per condition. **(i)** Quantification of the number of P bodies per cell in cells expressing ASAPΔtevs with indicated periods of light. N=20 cells per condition. Data are graphed as mean ± S.D. and are representative of 2 independent experiments. Statistical analyses were performed using a (**h**) One Way ANOVA or (**i**) a Student’s T-test.

**(j-k)** HEK293T cells transfected with **(j)** ASAP or **(k)** CTRL and treated with 450nm pulsed light for indicated periods of time followed by IF. Quantification of P body size. N=18-20 cells per condition. Data are graphed as mean ± S.D. and are representative of 2 independent experiments. Statistical analyses were performed using a One Way ANOVA.

**(l-n)** HCT116 cells transfected with ASAP and treated with 450nm pulsed light for indicated periods of time followed by IF. **(l)** Representative IF images of the P body marker, DCP1A, and Myc-tag for ASAP. Scale bar = 5μM. Quantification of (**m**) the number of P bodies per cell and **(n)** P body size. N=18-21 cells per condition. Data are graphed as mean ± S.D. and are representative of 3 independent experiments. Statistical analyses were performed using a One Way ANOVA.

**(o-q)** HCT116 cells transfected with CTRL and treated with 450nm pulsed light for indicated periods of time followed by IF. **(o)** Representative IF images of the P body marker, DCP1A, and Myc-tag for ASAP. Scale bar = 5μM. Quantification of (**p**) the number of P bodies per cell and **(q)** and P body size. N=19-20 cells per condition. Data are graphed as mean ± S.D. and are representative of 3 independent experiments. Statistical analyses were performed using a One Way ANOVA.

**(r-s)** HEK293T cells transfected with ASAP and treated with 450nm pulsed light for indicated periods of time followed by IF. **(r)** Representative IF images of the stress granule marker, G3BP, and Myc-tag for ASAP. Scale bar = 5μM. **(s)** Quantification in stress granule count per cell. N=21-23 cells per condition. Data are graphed as mean ± S.D. and are representative of 2 independent experiments. Statistical analyses were performed using a One Way ANOVA.

**(t-u)** HEK293T cells transfected with CTRL and treated with 450nm pulsed light for indicated periods of time followed by IF. **(t)** Representative IF images of the stress granule marker, G3BP, and Myc-tag for ASAP. Scale bar = 5μM. **(u)** Quantification in stress granule count per cell. N=21 cells per condition. Data are graphed as mean ± S.D. and are representative of 2 independent experiments. Statistical analyses were performed using a One Way ANOVA.

**Extended Data Fig. 7: Nucleoporins accumulate in P bodies after autophagy inhibition via NUP358. Related to Figure 6.**

**(a-c)** HEK293T cells transfected with CTRL or ASAP and treated with 450nm pulsed light for indicated periods of time followed by IF. **(a)** Representative IF images of DCP1A, MAb414, and Myc-tag for ASAP in ASAP expressing cells. **(b)** Representative IF images of DCP1A, MAb414, and Myc-tag for CTRL in CTL expressing cells. Scale bar = 5μM. **(c)** Quantification of the percent of P bodies colocalized with MAb414 in CTL expressing cells. N=24-29 cells per condition.

(**d**) HEK293T cells transfected with ASAP and treated with 450nm pulsed light for indicated periods of time followed by IF. Quantification of the percent of P bodies that colocalizes with cytoplasmic NUP358 signal, representative images shown in Fig. 6c. Data are graphed as mean ± S.D. and are representative of 2 independent experiments. Statistical analyses were performed using a One Way ANOVA.

(**e**) HEK293T cells transfected with ASAP and treated with 450nm pulsed light for indicated periods of time followed by IF. Quantification of the percent of cytoplasmic MAb414 signal that colocalizes with P bodies, representative images shown in Extended Data Fig. 7a. N=14-22 cells per condition. Data are graphed as mean ± S.D. and are representative of 2 independent experiments. Statistical analyses were performed using a One Way ANOVA.

(**f-g**) HEK293T cells transfected with ASAP and treated with 450nm pulsed light for 1hr followed by IF. **(f)** Representative IF images DCP1A, the ER marker SERCA2, and Myc-tag for ASAP. Scale bar = 5μM. **(g**) Quantification of the percent of P bodies that colocalizes with SERCA2 relative to the total number of P bodies. N=21-22 cells per condition. Data are graphed as mean ± S.D. and are representative of 2 independent experiments. Statistical analyses were performed using a Student’s T-test.

(**h-i**) HEK293T cells with stable expression of shScramble (shScr) or shNUP358. (**h**) qRT-PCR for gene expression of NUP358, data graphed as the mean ± S.D and are representative of 2 independent experiments. (**i**) Quantification of the NUP358 signal that is cytoplasmic or on the nuclear rim, data graphed as fold change to the respective dark condition for cytoplasmic and rim signal. N=19-23 cells per condition. Data are graphed as mean ± S.D. and are representative of 2 independent experiments. Statistical analyses were performed using a One Way ANOVA.

**Extended Data Figure 8: Light box and set up used for optogenetic manipulation of ASAP. Related to methods.**

(**a-f**) Amino acid sequences for all ASAP and CTRL constructs, highlighted according the corresponding region of construct indicated below each sequence.

(**g**) Images of light box fitted with LED light strips.

(**h**) Images of light box connected to light timer with lights off (left) and on (right).

**(i-j)** Images of light box in a standard tissue culture incubator **(i)** under ambient light and **(j)** using overhead red lights for optimal working conditions when harvesting samples.

**Extended Data Figure 9: Uncropped western blots.**

Corresponding figure and antibody are indicated. The dotted box represents where the blot was cropped for the figure.
